# Glycan-coated nanoparticles mimicking the ischemic glycocalyx scavenge the complement system conferring protection after experimental ischemic stroke

**DOI:** 10.64898/2026.03.30.715069

**Authors:** Gizem Mansour, Serena Seminara, Domenico Mercurio, Aurora Bianchi, Alessia Porta, Chantal Dembech, Patricia Perez Schmidt, Laura Polito, Claudia Durall, Franca Orsini, Luana Fioriti, Davide Comolli, Massimiliano De Paola, Gianluigi Forloni, Maria-Grazia De Simoni, Marco Gobbi, Stefano Fumagalli

## Abstract

Glycoproteins lining the luminal endothelial surface form the glycocalyx, composing the tripartite blood brain barrier. We explored the glycocalyx as a source of danger signals for complement lectin pathway after ischemic stroke. Our data indicate that hypoxic microvascular cells increased α-D-mannosyl and N-acetylglucosaminyl exposure after re-oxygenation, favoring mannose binding lectin (MBL) pathogenic deposition, and overexpression of inflammatory genes (*ICAM-1* and *MMP-2*). The hypoxia-conditioned medium induced neuronal damage (reduced MAP-2), microglia and astrocytic reactivity (increased/thickened ramifications) when applied to induced pluripotent stem cell-derived neurons, astrocytes and microglia co-cultures. All these effects were counteracted by mannose-capped gold nanoparticles (Man-GNPs), shown to bind and sequester MBL from the medium. We then tested the Man-GNPs *in vivo*, in an ischemic stroke model using humanized mice, knocked-in for human MBL. The ischemic mice (males:females 1:1) treated with Man-GNPs (3h after the ischemic onset) exhibited less anxiety at the elevated plus maze and reduced neuronal loss at 8d after ischemia compared to vehicle-treated. Thus, multivalent Man-GNPs represent a promising approach to take MBL away from its glycoproteic targets on the ischemic endothelium, hence preventing downstream pathogenesis. Moreover, these data support circulating MBL as a druggable pharmacological target to prevent the thrombo-inflammatory events following acute brain injury.

## Background

Glycans are sugar-based polymers, composed of many monosaccharides linked by glycosidic bonds which forms chain-like structures on cells and a large proportion of proteins^1^. They are implicated in a number of key physiological and pathological processes, including detection of infectious agents, cell adhesion, receptor activation, signal transduction and endocytosis^2^. Glycans like mannose, N-acetylglucosamine, glucose, fucose, sialic acid, and heparan sulfate are present in the glycoproteins lining the luminal surface of blood vessels, forming the glycocalyx. This structure physiologically supports vessel tone and integrity, provides mechanotransducted signals, interacts with cytokines and growth factors and regulates immune cell adhesion and rolling^3,4^. In the brain vasculature, the glycocalyx contributes to the blood brain barrier (BBB), as demonstrated by Kutuzov et al.^5^, who proposed the notion of a tripartite BBB, a sequence of diffusional constraints represented – starting from the luminal side - by the glycocalyx, the endothelium and the extravascular components.

As carbohydrate binding proteins, lectins are the major interacting partners of the glycans. Lectins have different affinity to varieties of sugars carrying mono-, di-, or multivalent carbohydrate binding sites^6–8^, and can either be free or exposed on the cell surfaces. Lectins are found in microbes^9–11^, plants^12–14^, animals including humans^15–19^, and are involved in various biological processes, such as cellular development, cell-to-cell signaling and immune responses. Therefore, the interaction between lectins and glycans plays a fundamental role in many pathophysiological processes. In humans, over 20 distinct types of lectins were detected including selectins, galectins and lectin pathway initiators^15,18^. The lectin pathway (LP) is part of the complement system, an arm of innate immunity triggered when specific lectin proteins, i.e. mannose binding lectin (MBL) or ficolin, bind to specific carbohydrates - and carbohydrate patterns - on the surface of pathogens (pathogen-associated molecular patterns, PAMPs) or expressed by damaged self-cells (damage-associated molecular patterns, DAMPs)^20^. While the complement activation downstream to the lectin binding to PAMPs has a protective role, that following the binding of DAMPs may be deleterious^21^. Among all the lectins, MBL shows a key pathogenic role after myocardial^22,23^, renal^24^, gastrointestinal^25^ and cerebral ischemic injury^26–29^. In all these conditions, but especially for cerebral ischemic injury (stroke), MBL appears to contribute to tissue injury, as supported by the fact that genetic deletion or pharmacological inhibition of MBL are protective^26,27,30^.

Stroke is the third leading cause of disabilities and second leading cause of death worldwide^31–34^. Among the different types of strokes, the ischemic one is the most frequent (88% of all strokes) and can be treated by the use of a thrombolytic agent and endovascular thrombectomy. Despite recent progress in prevention and management, stroke remains a medical issue considering that the available therapies present limitations like a short therapeutic window, neurotoxicity and increased risk of hemorrhages^35^. As such, around 40% of ischemic stroke patients are untreated. Those receiving the therapies may still show a limited improvement of stroke outcomes, even if a successful reperfusion is achieved, the so-called futile reperfusion^36^. Also, secondary events in the days that follows acute ischemic stroke cause vascular dysfunctions, due to thrombo-inflammation^37^ and no-reflow^38^, contributing to pathological processes. MBL is a hub of thrombo-inflammatory events^30,39^, hence representing a pharmacological target, whose druggability is warranted by the fact that it is a circulating protein, selectively released by the liver.

Considering that MBL has an oligomeric structure with multiple carbohydrate recognition domains (CRD)^40,41^, inhibitory compounds targeting it should present some degree of multivalency. Nanoparticles are ideal candidates offering multivalent binding of MBL if properly functionalized. In this study, glycan-capped gold nanoparticles (glyco-GNPs), i.e. mannose-capped gold nanoparticles (Man-GNPs) and glucose-capped gold nanoparticles (Glc-GNPs) were tested on experimental models of ischemic stroke. These GNPs were selected after an in vitro screening assessing their ability to inhibit MBL by a specifically-developed surface plasmon resonance assay, previously reported^42^.

While the complement system is an evolution-preserved pre-antibody inflammatory response, some key differences exist among its homologs in different species. This is the case of MBL, which has two isoforms (MBL-A and -C) in rodents – where most of preclinical studies were conducted on – and one functional isoform in humans (hMBL)^43^. To add translational value to our work, we explored the inhibition of hMBL in an in vitro model of ischemia with human-derived cells and in an in vivo model with a humanized mouse strain, i.e. knocked-out of murine MBLs and knocked-in of hMBL. In these mice, multivalent glyco-GNPs provided a modest, but significant reduction of behavioral deficits and neuronal loss when injected as a single bolus at 3h after the ischemic onset. Our findings indicate the therapeutic potential of glyco-GNPs, although further optimization of pharmacokinetic and treatment schedule is required to improve ischemic protection.

## Methods

### In vitro and in vivo models

#### In vitro model of hypoxia/re-oxygenation on endothelial cells

Immortalized human brain microvascular endothelial cells (ihBMECs) (5000 cells/cm^2^) (Innoprot) were seeded on black 96-well µ-plates (ibidi, Germany) with optically clear flat bottom, suitable for fluorescence microscopy. Plates were coated overnight before use, with fibronectin (Sigma) 15 mg/mL in Dulbecco’s phosphate buffer saline (DPBS, Euroclone). Cells were cultured in MCDB-131 (Gibco) supplemented with 5% fetal bovine serum (FBS, Euroclone), 2 mM L-glutamine (Gibco), penicillin/streptomycin (P: 100 U/mL-S: 100 U/mL, Sigma), 1 µg/mL hydrocortisone (Sigma), 50 µg/mL endothelial cell growth supplement (ECGS, Sigma), and kept at 37℃, with 5% CO_2_ and 90% humidity. Once cells formed monolayer, they were transferred into a hypoxic chamber (Whitley H35 Hypoxystation, Don Whitley Scientific, UK) at 37°C, and maintained in deoxygenated culture medium at the following gas concentrations: O_2_ 0.5%, CO_2_ 5% and N_2_ 94.5% for 16 h. Deoxygenated culture medium was obtained by leaving growth medium in a Petri dish inside the hypoxic chamber for 7 h before use. Control cells were maintained in fresh culture medium in a normoxic incubator. After 16 h, hypoxic cells were taken out of the hypoxic chamber, washed twice with DPBS and re-oxygenated for 4 h in a normoxic incubator. During re-oxygenation, normoxic and hypoxic cells were exposed to 30% human serum (HS, Innovative Research) with or without sugar-GNPs (40-20-5 µg/mL) or MBL depleted human serum 30% (MBL Dep. HS 30%). All serums were diluted in culture medium (not supplemented with FBS). At the end of re-oxygenation, media of all conditions collected and immediately stored at −20°C. Cells were either fixed with 4% paraformaldehyde (PFA) in DPBS RT for 15 min or scratch and collected with lysis buffer (PureLink RNA miniKit, Invitrogen, CA, USA) containing 1% of 2-mercaptoethanol. Fixed cells were stored in 0.01M NaN_3_ in PBS 0.01M at 4°C until immunofluorescence analysis, collected cells were stored at 20°C until RT-PCR analysis.

#### hIPSC-derived co-cultures of neurons, astrocytes and microglia

Episomal hIPSC from healthy donor were obtained from Gibco™ (Life Technologies, CA, US, Lot V2.0). The hIPSC line was cultured and expanded in feeder-free conditions by passaging every 3-5 days when they reached 70%-80% confluence, in a xeno-free cell culture medium formulation (StemMACS™ iPS-Brew XF, Miltenyi Biotec S.r.l.). The cells were differentiated towards the neuronal lineage by blocking the TGF-β/BMP-dependant SMAD signalling (STEMdiff™ SMADi Induction Kit, STEMCELLS™ Technologies), resulting in efficient neural induction. By day 7 of neural induction, primitive neuronal progenitors were dissociated with Accutase (Life Technologies), passed through a 100-μm strainer, and plated on Matrigel-coated dishes in an expansion medium (STEMdiff™ Neural Progenitor Medium, STEMCELLS™ Technologies). Astrocyte cultures were obtained from NPCs at the sixth to the tenth passage (P6-10) by applying specific culture conditions able to generate differentiated and functionally active neurons and astrocytes^44,45^ in Astrocytes Medium (ScienCell Research Laboratories, Inc.). For microglia production, iPSCs were first differentiated toward a mesodermal, hematopoietic lineage with, then highly pure populations of non-adherent CD43+ hematopoietic progenitors were differentiated in homeostatic microglia with the addition of specific microglia differentiation and maturation kits (STEMdiff™, STEMCELLS™ Technologies)^46^. After 47 days in culture, mature microglia were collected and added to the cells differentiated to the neuronal lineage to establish the co-cultures. Mixed neuron/astrocyte/microglia co-cultures will be obtained by inducing neuron differentiation to NPCs seeded on a mature astrocyte layer and then adding differentiated microglia to the co-culture one week before drug testing or analysis.

#### In vivo model of ischemic stroke

Procedures involving animals and their care were conducted in conformity with institutional guidelines in compliance with national and international laws and policies. Our project involving animals received ethical approval by the local ethic committee and by the Italian Ministry of Health (Autorizzazione n° 383/2021-PR, project number 9F5F5.194). Male and female C57BL/6 J mice were used at 10-13 weeks old and weighting 26–28 g, either WT or hMBL KI, kindly provided by Dr. Gregory Stahl, Harvard Medical School, Boston, US, and colonized at the Mario Negri Institute). The protocols and details of this report are in accordance with the ARRIVE guidelines ( http://www.nc3rs.org.uk/page.asp?id=1357, see the list provided as a supplementary file).

Focal cerebral ischemia was induced by transient middle cerebral artery occlusion (tMCAo) as previously reported by our group^43^. Anesthesia was induced by 3% and maintained by 1.5% isoflurane inhalation in an N_2_O/O_2_ (70/30%) mixture. Transient ischemia was achieved using the filament model. A silicone coated monofilament nylon suture (sized 7-0, diameter 0.06-0.09 mm, length 20 mm; diameter with coating 0.23 mm; coating length 10±1 mm, Doccol Corporation, Redlands, CA, USA) was introduced into the common carotid artery (appropriately isolated) and advanced to block the middle cerebral artery (MCA). After 30 minutes, blood flow was restored by carefully removing the filament. During surgery body temperature was kept at 37°C by a heating pad. During MCA occlusion, mice were awakened from anesthesia, kept in a warm box and tested for intraischemic deficits (for inclusion/exclusion criteria, see below). Analgesia was achieved by local application of an ointment (EMLA, containing 2.5% lidocaine and prilocaine, Aspen Pharma) where the skin was opened. Animals were monitored after surgery according to the IMPROVE guidelines^47^ and treated subcutaneously with 0.1 mg/kg buprenorphine if developing severe signs of distress (IMPROVE’s amber category of clinical signs).

##### Inclusion/exclusion criteria

Mice were included in the study if successfully induced with ischemia, i.e., filament correctly positioned in the MCA. Thus, ischemic animals were included if presenting ≥3 of the following intra-ischemic deficits:

1. the palpebral fissure had an ellipsoidal shape (not the normal circular one)
2. one or both ears extended laterally
3. asymmetric body bending on the ischemic side
4. limbs extended laterally and did not align to the body.

Mice were excluded if:

1. they died during MCA surgery
2. they had a decrease in body weight < 35% before sacrifice compared to baseline

### Preparation of glyco-GNPs

Gold nanoparticles (GNPs) were synthesized and characterized as previously described^42,48^. Briefly, sodium citrate (9 mL, 68 mM), HAuCl_4_ (7.5 mL, 10 mM) and AgNO_3_ (490 μL, 5.9 mM) were mixed and stirred for 6 min at room temperature. Then the solution was added into 250 mL of boiling water in a 500 mL flask and stirred (750 rpm) for 1 h at 100°C, allowing the formation of the seeds. The seed solution was cooled to room temperature, 5 mL of glycerol was added, and the solution was left stirring for 10 min. A second mixture of sodium citrate (10 mL, 34 mM), HAuCl_4_ (7.5 mL, 10 mM) and AgNO_3_ (426 μL, 5.9 mM) was pre-mixed for 6 min and then added to the seed solution, followed immediately by the addition of hydroquinone (8 mL, 91 mM). The solution was left stirring at 750 rpm for 1 h to age and improve homogeneity. Synthesized citrate-capped GNPs were purified by centrifuging, using Millipore Amicon Ultra-4 Centrifugal Filter Units, 30KDa cut-off, at 6000 rpm for 4 min and stored as colloidal solution in water (12 mL, Au content 14 mg). GNPs were functionalized by capping the metal surface with thiol-glyco-derivatives, exploiting the strong thiol-gold bond and allowing a self-assembled monolayer on the gold surface^42^.

### Quartz Crystal Microbalance (QCM)

Attana CellTM 200 QCM was used to investigate possible hypoxia related changes on glycocalyx of ihBMECs. For that, the LNB-cell-optimized polystyrene (LNB-COP) surface was coated with fibronectin (15 mg/mL) overnight at 37°C. The ihBMECs, suspended in 100 µL of supplemented MCDB-131 culture medium, were seeded onto the LNB-COP surface at a density of 100’000/cm^2^, and incubated for 48 h at 37 °C in 5% CO_2_ to achieve a monolayer distribution. Then, the cells underwent in vitro hypoxia/re-oxygenation described above. At the experimental endpoint, ihBMEC cells were washed with PBS and fixed with cold 4% paraformaldehyde for 15 min at RT. The CTRL and HYPX chip pairs were inserted in a CellTM 200 QCM instrument and the flow rate was set at 10 µL/min at 22C°. The kinetic binding experiments were performed when the baseline drift signal was ≤0.2 Hz/min A blank injection was performed before each analyte injection, and it was used to correct the baseline drift. Four different concentrations (0.7-2-6-18 µg/mL) of the plant lectins Concanavalin-A (ConA) and the wheat germ agglutin (WGA) with sugar affinities for α-D-mannosyl and N-acetylglucosaminyl, respectively, were injected for 250s at 10 μL/min over CTRL and HYPX ihBMEC on LNB-COP chips with a 600s dissociation. The running and dilution buffers were TBST (10 mM Tris buffer [pH 7.4], with 150 mM NaCl and 0.005% Tween-20) containing 1.2 mM CaCl_2_. Regeneration of the chip was done after each injection of lectins with TBST buffer without Ca2+ and two glycine [pH 1.5] plug volume 5.0 injections, respectively. Cell coverage and MBL deposition were then determined by Hoechst staining and visualizing CTRL and HYPX ihBMECs on LNB-COP under a fluorescent microscope. The data was recorded using the AttesterTM software (Attana AB) and the data was prepared and analyzed using the EvaluationTM (Attana AB).

### Protein corona evaluation

Protein corona was formed by 1h of incubation of sugar-GNPs (80 µg/mL) in 30% HS in TBST/Ca^+2^. The proteins associated with the “soft” corona (i.e., those with a relatively low affinity) were progressively removed by four consequent washing – centrifugation steps in TBST/Ca^+2^ at 13 rcf, 40 min ^49^. Samples were immediately stored at −20 °C. Presence of MBL in the soft and hard corona was revealed with western blot analysis. Equal volumes of protein corona samples (10 µl/sample) were electrophoresed on a 12% sodium dodecyl sulfate-polyacrylamide gel under denaturing conditions, using a mixture containing 2% SDS and heating the samples at 96 °C for 5 minutes. The proteins were then transferred to polyvinylidene fluoride membrane. Anti-hMBL mouse antibody (clone 3E7) (1:1000; Hycult Biotech) followed by anti-mouse HRP-conjugated antibodies 1:10000, goat anti-mouse IgG specific light chain, Jackson ImmunoResearch) were used. Immunocomplexes were visualized by chemiluminescence using the IMMOBILON western blot substrate (Merck KGaA).

### Immunofluorescence

#### In vitro model

Immunofluorescence was performed on formaldehyde fixed cells. Nuclei were stained with DAPI (1 mg/mL, Invitrogen). For MBL staining, after a blockade with 1% normal goat serum for 1 h, fixed cells were incubated overnight with mouse antihuman MBL (clone 3E7, 1:100, Hycult Biotechnology). Cells were then incubated with biotinylated anti-mouse (1:200) for 1 h followed by incubation with streptavidin Alexa 647-conjugated (1:100) for 30 min. For F-actin staining, fixed cells were blocked with 1% bovine serum albumin for 30 min, then incubated with phalloidin Alexa 488-conjugated (1:100, Invitrogen).

Immunofluorescence on hIPSC-derived cultures was done on formaldehyde fixed cells. For neuronal and astrocytes analysis, cells were permeabilized for 30 min with a solution containing 0.4% triton, 5% fetal bovine serum in DPBS, then incubated overnight at 4°C with antibodies against MAP-2 (rabbit, 1:500, Merck Millipore), β3-Tubulin (mouse, 1:200, BioLegend), Iba-1 (rabbit, 1:200, FUJIFILM Wako Pure Chemical Corporation) and GFAP (mouse, 1:500, Merck Millipore). Cells were then incubated with secondary antibodies conjugated with Alexa fluor488, −555, −594 or −640 (1:1000, Invitrogen) for 2 h and nuclei were counterstained with Hoechst for 30 min. Then coverslip glasses were mounted on microscope slides using ProLong™ Gold Antifade Mountant (Invitrogen) before images acquisition.

#### In vivo model

For MBL deposition, 20 µm-thick coronal sections were incubated overnight with mouse anti-human MBL (1:100, Hycult Biotechnology). Alexa546 fluoro-conjugated goat-anti-rat IgG (1:500, Molecular Probes) was used as a secondary antibody. Brain vessels and nuclei were stained with Alexa488 fluor-conjugated isolectin (IB4, 1:200, Invitrogen) and Hoechst (1μg/ml, Invitrogen), respectively. For glycoprofiling, the same sections of mouse brains were incubated for 2h with Alexa546 fluor-conjugated with fluorescent plant lectins as detailed in ^50^.

### Confocal Microscopy

#### In vitro models

Acquisitions were done by microscopy with a confocal scanning A1 unit (Nikon), managed by ‘NIS-elements’ software. We used a sequential scanning mode to avoid bleed-through effects. Graphic elaboration of images was done with GIMP software.

Images for MBL deposition on normoxic and hypoxic ihBMECs treated with glyco-GNPs, were obtained by acquiring a 2.5 mm field in the center of each well. Cells were illuminated with 405 nm (nuclei) and 640 nm (MBL) lasers; 512 pixel images were acquired with a 20x objective, over a 10 µm stack, with 2.65 µm step size, and stitched with 15% overlay. After background subtraction, the fluorescent signal for MBL was quantified by ImageJ software and expressed as integrated density.

Images for the quantification of neurons, astrocytes and microglia in the hIPSC-derived co-cultures were obtained by acquiring a 1.2 mm field in the center of each well. Cells were illuminated with 405 nm (nuclei), 488 nm (GFAP), 561 (MAP-2) and 640 nm (Iba1) lasers; 1024 pixel images were acquired with a 40x objective, over a 12 µm stack, with 0.98 µm step size and stitched with 15% overlay. Eight, non-overlaid regions of interest sized 25×250x12 µm were stereologically sampled from the overview images to proceed with fluorescent signal measures. For MAP-2 and GFAP analysis, we segmented the positive signal using the isosurface function of Imaris and calculated the total occupied volume expressed as µm^3^. As an additional GFAP analysis, we identified the first branches arising from the soma of randomly selected cells and traced a line crossing the branch perpendicularly. GFAP signal intensity (gray level) over the traced line served to calculate branch thickness using the full-width-at-half-maximum (FWHM) method using ImageJ. For Iba1 analysis, positive cells were isolated, segmented and skeletonized using an originally developed ImageJ macro. The skeletonized objects were analyzed using the 2D segmentation plug-in of ImageJ to compute number and length of branches, number of junctions (branch connecting nodes).

#### In vivo model

Images were acquired by confocal microscopy using a scanning sequential mode to avoid bleed-through effects by an IX81 microscope equipped with a confocal scan unit FV500 with 3 laser lines: 405 nm (nuclei), 488 nm (vessels) and 546 nm (MBL) lasers and a UV diode. Three-dimensional images were acquired over a 10 μm z-axis with a 0.23 μm step size and processed using Imaris software (Bitplane) and GIMP software.

For the quantification of the plant lectin signal, confocal microscopy was done using a sequential scanning mode to avoid bleed-through effects by an A1 Nikon confocal microscope with an excitation at 405 nm for DAPI, 488 for WGA, 561 nm for ConA and 640 for IB4 signals. Large view images were acquired by a 20x 0.5 NA objective, with pixel size 0.62 μm and automatically stitched with 10% overlap. Large view images served as reference to identify the cortical regions of interest for subsequent analysis. Four three-dimensional volumes sized 210 x 210 x 12 µm were acquired with a 40x 0.75 NA objective. Digital image analysis was done using originally developed ImageJ plugins. Briefly, the image was corrected for background noise and then the raw integrated density of pixels calculated over all focal planes. The sum of focal planes’ raw integrated density was used for statistical analysis.

### Reflectance Confocal Microscopy

Reflectance confocal microscopy was set up to visualize glyco-GNPs. Man-GNPs, Glc-GNPs and Alexa488-conjugated beads were dropped on different microscope slides. Focus and background adjustment were illuminated with 488 nm (Alexa488-conjugated beads) lasers and glyco-GNPs were excited with 561 nm (HV: 100, laser power: 0.5) changing the dichroic to the transmitted/reflectance mirror (BS20/80) with pinhole adjustment. Images were acquired with a 100x 1.49 NA oil immersion objective. Same setting then applied on normoxic and hypoxic ihBMECs treated with 30% HS in presence of Man-GNPs to visualize nanoparticles. Images were obtained by acquiring a 2.5 mm field in the center of each well.

### Structured Illumination Microscopy (SIM)

Structured illumination microscopy (SIM) was done on a Nikon SIM system with a 100x 1.49 NA oil immersion objective (for F-actin quantification), managed by NIS elements software. Cells were imaged at laser excitation of 405 for nuclei, 488 for F-actin with a 3D-SIM acquisition protocol. Fourteen-bit images sized 1024 pixels with a single pixel of 0.030 mm (100) or 0.036 mm (60) were acquired in a gray level range of 0–4000 to exploit the linear range of the camera (iXon ultra DU-897U, Andor) at 14-bit and to avoid saturation. Raw and reconstructed images were analyzed with the SIM check plugin of ImageJ. For cytoskeletal organization, 12 cells from four wells per condition were acquired and analyzed with F-actin touches on 1 µm-spaced grid superimposition. SIM images were quantified with ImageJ. Briefly, a region of interest was drawn including the cell cytoplasm but not the nucleus. Background noise was normalized throughout the samples and F-actin filaments were selected by signal segmentation followed by the skeletonize function.

### Real time RT PCR

Total RNA was extracted from cells at the end of the 4 h re-oxygenation using the PureLink RNA miniKit (Invitrogen, CA, USA) according to the manufacturer’s instructions. Samples of total RNA (100 ng) were treated with DNAse (Applied Biosystems, Foster City, CA, USA) and reverse-transcribed with random hexamer primers using multiscribe reverse transcriptase (TaqMan reverse transcription reagents, Applied Biosystems, Foster City, CA, USA). Primers were designed to span exon junctions in order to amplify only spliced RNA, using PRIMER-3 software (http://frodo.wi.mit.edu/) based on GenBank accession numbers (ß -actin: NM_001101.5, MBL2: NM_000242.2). The same starting concentrations of cDNA template were used in all cases. Real-time PCR was done using Power SYBR Green according to the manufacturer’s instructions (Applied Biosystems). ß-actin was used as reference gene and relative gene expression levels were determined according to the Ct method (Applied Biosystems). Data are presented with individual values. Primer sequences: IL-1α fwd: tgaagaagacagttcctccattg, rev: cttcatggagtgggccatag; MMP-2 fwd: atgccgcctttaactggag rev: ggaagccaggatccattttc; ICAM-1 fwd: tgatgggcagtcaacagcta rev: ggtaaggttcttgcccactgg; β-actin fwd: ccagctcaccatggatgatg, rev: atgccggagccgttgtc.

### Treatments

Mice received by intravenous (IV) bolus injection in the tail vein at 3h after onset of tMCAo, while under anesthesia, 20 μg/mL (40 μg/mouse) Man-GNPs and Glc-GNPs (injected volumes maximally 100 μL/mouse). Vehicle treatment was done by buffer injections. Twelve mice for vehicle and Man-GNPs and eleven mice for Glc-GNPs treatment were used with 1:1 gender ratio. Mortality at the experimental endpoint (8d) was similar for all groups (25%).

### Plasma collection

Blood (≈50 μL) was obtained from the submandibular vein (cheek pouch) -2d, 90’ and 48h after tMCAo. Drops of blood were exude from the stick point and were collected into a suitable sample container. Blood was centrifuged at 2000 g for 15 minutes at 4°C and the plasma was stored at −80 °C for subsequent analyses.

### Western blotting

Plasma samples from hMBL KI mice at -2d, 90’ and 48h after tMCAo were collected and immediately stored at −80°C. Equal amounts of plasma proteins (10 µg/sample) were electrophoresed on Tris-Acetate 3-8% gradient gels (EA3735, Invitrogen) in 1% SDS loading buffer using Tricine-SDS running buffer (Novex LC1675, Invitrogen) and transferred to nitrocellulose membranes by TransBlotTurbo system (Bio-rad). Membranes were incubated overnight at 4°C with a mix of hMBL mouse antibody (clone 3E7) (1_1000; Hycult Biotech) and hMASP-2 rat antibody 1:1000 (clone 8B5). Membranes were then incubated for 1h at RT with a-mouse IRDye800 (1:10000, LI-CORE Bioscience) together with an a-rat Alexa 647 (1:5000, Invitrogen), used for hMBL and MASP-2 staining respectively. Immunocomplexes were visualized by fluorescence using ChemiDoc Imaging Systems (Bio-Rad). Quantifications were done with Image Lab Software (Bio-Rad) and results were standardized using the total protein loaded (Ponceau solution).

### Behavioral tests

#### Neuroscore

Each mouse was rated on two neurologic function scales. The general deficit scale evaluates hair (0–2), ears (0–2), eyes (0–4), posture (0–4), spontaneous activity (0–4), and epileptic behavior (0–12), whereas the focal deficit scale evaluates body symmetry (0–4), gait (0–4), climbing on a surface held at 45° (0–4), circling behavior (0–4), front limb symmetry (0–4), compulsory circling (0–4), and whisker response to a light touch (0–4). Scores range from 0 (healthy) to 56 (the worst performance in all categories) and represent the sum of the results of general and focal deficits (13 categories). Results are expressed as composite neurological score. All mice meeting the criteria (intraischemic deficits) were included. A trained investigator who was blinded to the experimental conditions performed experiments.

#### Elevated plus maze

The elevated plus maze (EPM) measures disinhibition and anxiety-like behaviours. The test consists of two open and two closed arms (each 35 cm × 5.5 cm), and a central platform (5.5 cm × 5.5 cm), elevated 60 cm above the ground. Mice were acclimatized in the room for 1 h prior to testing, then placed in the central platform facing an open arm and their movements were recorded for 5 min. Video recording and time spent in the closed and open arms were measured by Ethovision XT, 514.0 (Noldus Information Technology, Wageningen, The Netherlands). All mice meeting the criteria (intraischemic deficits) were included. Data were expressed as percentage of time spent on the open arms (OA) over the total time the mouse spent on the maze [open arm time % = 100 × (OA time/total time)].

### Brain collection

Mice were anesthetized with 300 µL of ketamine (150 mg/kg) plus medetomidine (2 mg/kg) intra-peritoneally before sacrifice. Mice were transcardially perfused with 30 mL of phosphate buffered saline (PBS) 0.1 mol/L, pH 7.4, followed by 60 mL of chilled paraformaldehyde (4%) in PBS. The brains were carefully removed from the skull and post-fixed for 6 h at 4 °C, then transferred to 30% sucrose in 0.1 mol/L phosphate buffer for 24 h until equilibration. The brains were frozen by immersion in isopentane at −45 °C for 3 min and stored at −80 °C until use.

### Neuronal count

The histological lesion at 7d after the ischemic onset was measured as the relative density of viable neurons in the ipsi-lateral striatum or cortex as compared to the contra-lateral. Three cresyl violet–stained 20 µm-thick coronal sections at −0.6, 0 and + 0.6 mm from the bregma were acquired from each mouse and visualized at 20x magnification with an Olympus BX61 Virtual Stage microscope, with a pixel size of 0.346 mm. Neuronal counting was performed by segmenting the cells in regions of interest designed in the cortex and striatum where the lesion was clearly detectable. The regions of interest on the contralateral side were positioned mirroring those on the ipsilateral side. A segmented circular signal smaller than the area threshold of 25 µm2, which is associated with glial cells was excluded from the analysis (supplementary figure 1). For quantification, we used ImageJ software ^51^.

### Statistics

Wells containing cells or animals were randomly allocated to treatments. Subsequent evaluations were done by blinded investigators. Data are reported as bar plots with mean ± SD. Groups were compared by analysis of variance (ANOVA) and post hoc test, as indicated in each figure legend. The parametric or non-parametric test was selected after a Kolmogorov–Smirnov test for normality to assess whether groups met normal distribution. The constancy of variances was checked by Bartlett’s test. Data with equal variances were analyzed with two-way ANOVA followed by Tukey’s and Sidak’s multiple comparison, one-way ANOVA followed by Tukey’s multiple comparisons or unpaired t-test. Welch’s corrected t-test was used for normally distributed data with unequal variances. Group size for in vitro studies was defined pre hoc using the formula: n = 2s^2^ f(a,b)/2 (sd in groups = s, type I error a = 0.01, type II error b = 0.2, percentage difference between groups = 200 (2 time fold-change in the hypoxic group)). The standard deviation between groups was calculated based on a pilot experiment presented in 36 measuring the overexpression of *ICAM-1* due to hypoxia, and resulted in s = 42, yielding n = 4).

Group size for in vivo studies was defined pre hoc using the formula: n = 2s^2^ f(a,b)/2 (sd in groups = s, type I error a = 0.05, type II error b = 0.2, percentage difference between groups = 25). The standard deviation between groups was calculated based on previous experiment with neuroscore at 48h as primary endpoint, and resulted in s = 22, yielding n = 12).

## Results

### Effects of hypoxia-reoxygenation on the glycocalyx of brain microvascular endothelial cells

Initial studies were carried out to characterize the hypoxia-related changes on the glycocalyx of ihBMECs, testing in particular the hypothesis of an increased exposure of MBL-targeted glycoproteins i.e., containing α-D-mannosyl and N-acetylglucosaminyl domains. For these analyses, we used quartz crystal microbalance (QCM), culturing brain microvascular endothelial cells on LNB-COP chips, which were then exposed in the QCM fluidic channels to different concentrations (0.7, 2, 6 and 18 µg/mL) of ConA (a plant lectin specific for α-D-mannosyl moieties) and WGA (a plant lectin specific for N-acetylglucosaminyl moieties). Cells were analyzed at the end of 16h hypoxia and at the end of 4h re-oxygenation in presence of human serum with or without MBL (experimental plan in Fig. 1A). Fig 1B reports the sensorgrams, showing that the binding of both ConA and WGA decreased at the end of hypoxia compared to normoxia, suggesting a damage of the glycocalyx. After re-oxygenation, regardless of MBL presence in the serum, both signals increased compared to normoxia (Fig. 1B), in line with an increased exposure of the α-D-mannosyl and N-acetylglucosaminyl, both targets of MBL. Consistently, MBL was indeed found deposited on hypoxic/re-oxygenated cells by confocal microscopy (Fig. 1C).

**Figure 1.**
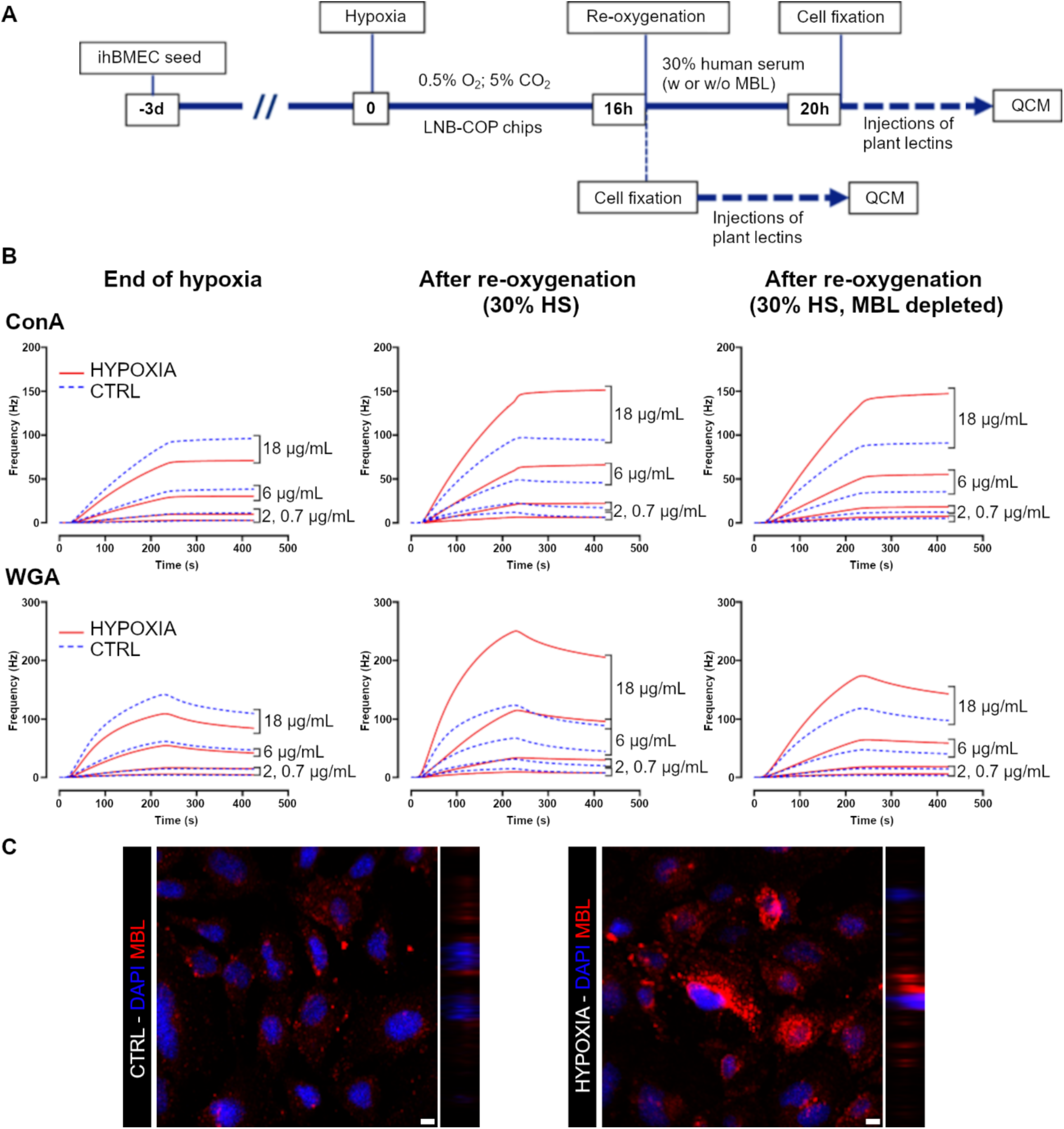
In vitro characterization of the brain microvascular glycocalyx in hypoxic conditions. **A**) The experimental plan for lectins binding to ihBMECs at the end of hypoxia or after re-oxygenation in absence/presence/. **B**) Sensorgrams obtained with Quartz Crystal Microbalance showing the binding of ConA (upper panels) and WGA (lower) injected at four different concentration (0.7-2-6-18 µg/mL) over chip-adherent ihBMECs. The data show decreased binding at the end of the 16h of hypoxia and increased binding after the 4h of re-oxygenation either in presence or absence of MBL compared to normoxic ihBMECs. **C**) Microphotographs of MBL (red) deposited on normoxic (left) or hypoxic (right) ihBMEC after re-oxygenation. Nuclei in blue (DAPI), scale bars 10 µm.

### MBL-binding properties of glycan-capped gold nanoparticles (GNPs) and their effects on hypoxic ihBMECs

We previously demonstrated with surface plasmon resonance (SPR) studies that Man-GNPs maintain their MBL-binding proteins in human serum, thus excluding an interfering effect of the protein corona formed on the nanoparticle^42^. To confirm and extend this finding, we used here a different approach evaluating, by Western blot, the presence of MBL in both the soft and the hard corona^52^ induced in Glyco-GNPs incubated in 30% HS diluted in TBST/Ca^+2 53^. The soft corona, composed of low affinity serum proteins, was progressively removed by four consequent washings/centrifugation steps in TBST/Ca^+2^, while the hard corona includes the high affinity proteins remaining bound to the nanoparticles after the washing. MBL was found on hard corona of Man-GNPs, confirming that protein corona formation did not interfere with hMBL targeting. This binding was strong and sugar specific as the signal detected on Glc-GNPs was at a lesser extent (Fig. 2A).

**Figure 2.**
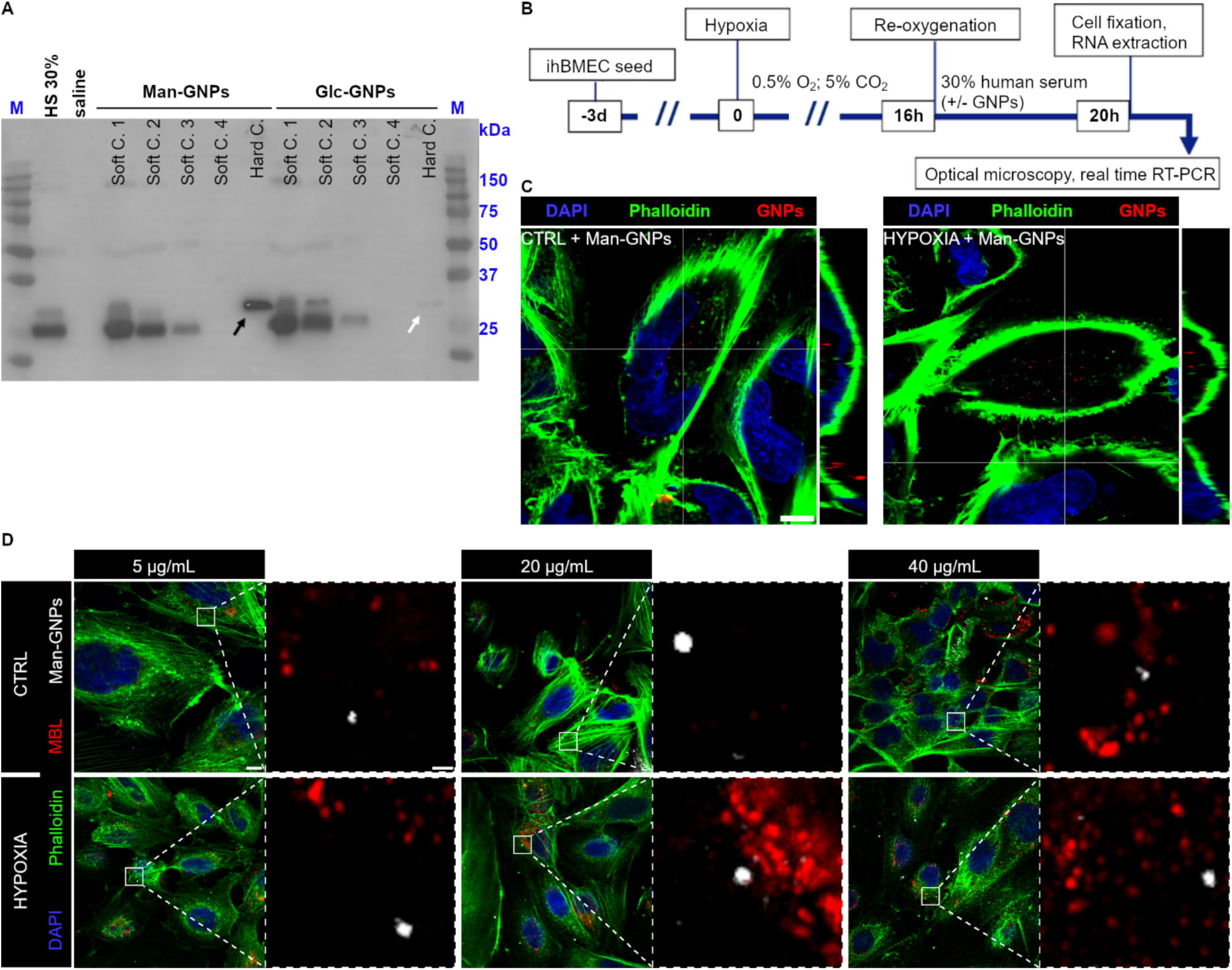
In vitro characterization of serum hMBL binding to GNPs under acellular conditions and its localization on cellular systems. **A**) MBL detection on the soft and hard corona samples obtained after preincubation of GNPs with human serum. The MBL signal decreased in the soft corona concomitantly with the three washes (Soft C1-3) and no signal was captured in the fourth wash (Soft C.4). The presence of MBL in the Hard Corona (Hard C., i.e. the proteins remaining after the washings because of their high affinity for the GNPs,) was strong for Man-GNPs (black arrow), and much less for Glc-GNPs (white arrow). **B**) The experimental plan for testing sugar-GNPs localization on ihBMEC. **C**) 3D microphotographs of Man-GNPs (red, reflectance microscopy) and F-actin (phalloidin, green) in normoxic (CTRL) or hypoxic (HYP) ihBMECs undergone re-oxygenation in the presence of 40 µg/mL Man-GNPs in 30% human serum. Man-GNPs were internalized in the cytoplasm of ihBMECs. Nuclei in blue (DAPI), scale bar 10 µm. **D**) Normoxic (CTRL) or hypoxic (HYPOXIA) ihBMECs undergone re-oxygenation in the presence of 5, 20 or 40 µg/mL Man-GNPs in 30% HS were analyzed by reflectance confocal microscopy for Man-GNPs and MBL co-localization. Microphotographs show that Man-GNPs (white, reflectance microscopy) and hMBL (red) did not co-localize (as seen in magnification of white frame, scale bar 1 µm). Phalloidin in green, nuclei in blue (DAPI), scale bar 10 µm.

To evaluate the effect of MBL-binding Man-GNPs on ihBMECs undergoing hypoxia/reoxygenation, different concentrations of Man-GNPs and Glc-GNPs (5, 20 and 40 µg/mL) were added to ihBMECs during the 4h re-oxygenation in presence of 30% HS (experimental plan in Fig. 2B). No toxic effects were observed even at the highest concentration (Supplementary Fig S1). GNPs could be visualized through reflectance confocal microscopy (RCM) allowing their unlabelled observation (Supplementary Fig. S2). In both normoxic and hypoxic ihBMECs, Man-GNPs were detected in the cell soma (Fig. 2C), and did not colocalized with deposited MBL (Fig 2D). Importantly, the presence of 5 and 20 µM Man-GNPs, during the re-oxygenation, almost completely antagonized the 2-fold increase of MBL deposition associated with hypoxia/re-oxygenation (Fig 3A-B). No effect was observed with 40 µM Man-GNPs.

**Figure 3.**
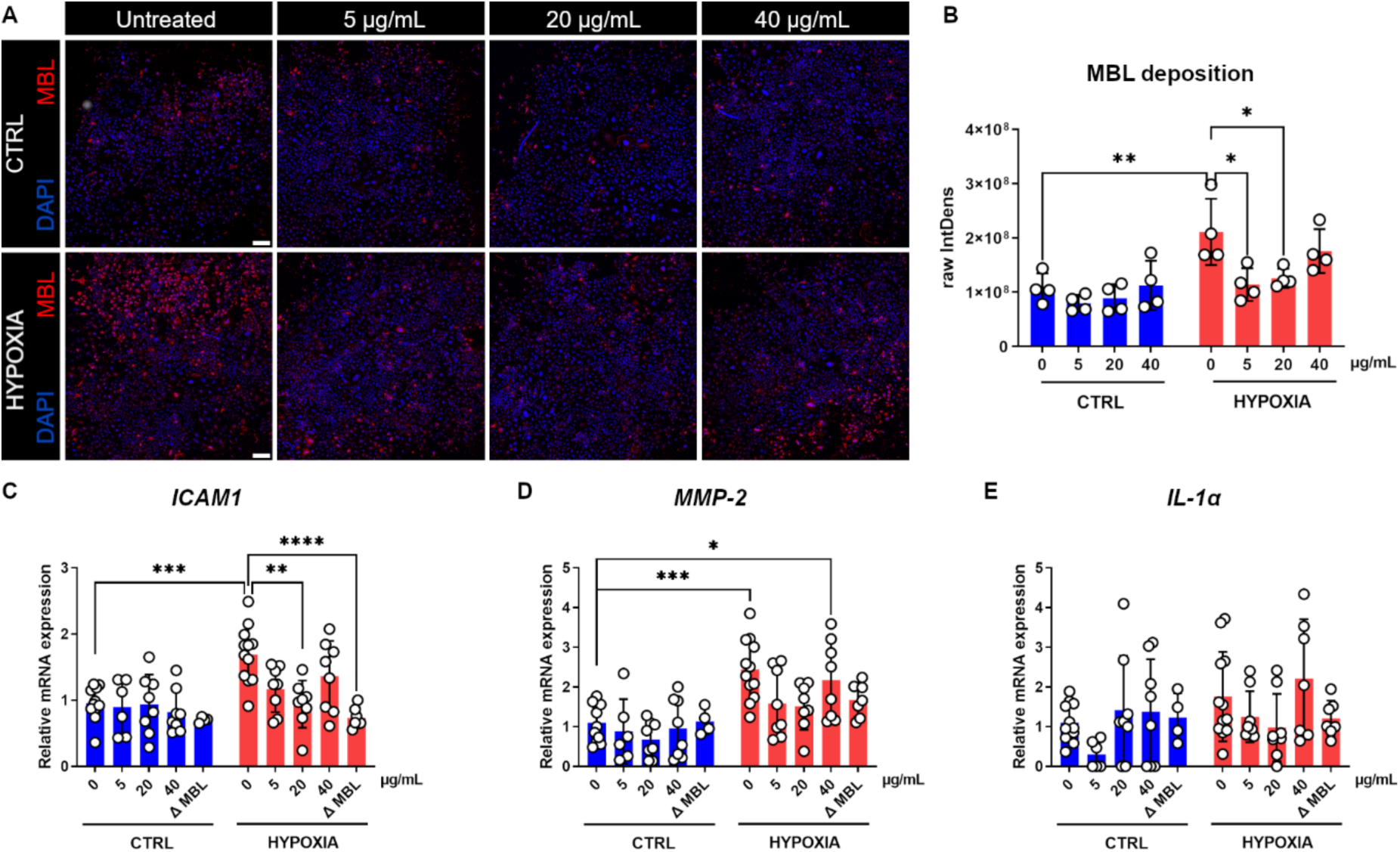
In vitro test of MBL inhibition by Man-GNPs, effects on microvascular inflammation. **A)** Microphotographs of MBL (red) deposited on normoxic (CTRL) or hypoxic (HYPOXIA) ihBMECs undergone re-oxygenation in the presence of 5, 20 or 40 µg/mL Man-GNPs in 30% HS (w/Man-GNPs). Nuclei in blue (DAPI), scale bar 200 µm. **B**) MBL deposition, measured as fluorescence intensity, was greater on hypoxic than normoxic cells exposed to 30% HS. This increase was significantly reduced when ihBMECs were exposed to 5 and 20 µg/mL of Man-GNPs. Data as mean with individual values ± SD (n= 4). Two-way ANOVA followed by Tukey’s multiple comparisons, **p<0.001, *p<0.05. **C**) Overexpression of *ICAM-1* in hypoxic ihBMECs was significantly reduced when the cells were exposed to 20 µg/mL of Man-GNPs, to a similar extent than exposure to MBL depleted HS 30% (Δ MBL). **D**) Overexpression of *MMP-2* in hypoxic ihBMECs was partially counteracted by 20 µg/mL of Man-GNPs. **E**) Expression of *IL-1α* was not significantly changed in presence of Man-GNPs with or without hypoxia. Data from 3 independent experiments, presented as mean with individual values ± SD (n= 4-12). Two-way ANOVA followed by Tukey’s multiple comparisons, ****p<0.0001, ***p<0.001, **p<0.01, *p<0.05.

Hypoxic ihBMECs upregulated the inflammatory markers *ICAM-1* (1.69 ± 0.4 fold-change, p<0.001), *MMP-*2 (2.45 ± 0.8, p<0.001) and *IL-1α* (1.76 ± 1.1, n.s.) compared to normoxic ihBMECs (Fig. 3D-F). The 20 µg/mL dosing of Man-GNPs reduced the overexpression of *ICAM-1* (to 0.94 ± 0.4 fold-change, p<0.01) and, at a lesser extent, *MMP-2* (1.52 ± 0.6 fold change) compared to untreated hypoxic ihBMECs. Similar effects were obtained re-oxygenating the ihBMECs with MBL-depleted 30% HS (Fig. 3C-E).

### Effects of reduced vascular inflammation by Man-GNPs on brain damage

We then sought to see if the reduction of hypoxia-induced vascular inflammation provided by Man-GNPs could translate in a lower damage of brain cells. To do so, conditioned medium (CM) of ihBMECs after 16h hypoxia and 4h re-oxygenation (HYP CM), with or without 20 µg/mL Man-GNPs, was collected, with normoxic medium used as control (NORM CM). The CMs were applied for 24h to co-cultures of human induced pluripotent stem cell (hIPSC)-derived neurons, astrocytes and microglia (experimental plan in Fig. 4A). HYP CM exposure induced the appearance of damaged neurons with a circular shape and no dendrites (arrows in Fig. 4B), which were less frequent if HYP CM were reoxygenated in the presence of Man-GNPs. Digital image analysis showed decreased neuronal MAP-2 stained volume in co-cultures exposed to HYP CM (6.0 ± 2.1 × 10^4^ µm^3^ ± SD), compared to NORM CM (11.1^4^ ± 2.4 × 10^4^), significantly prevented in co-cultures exposed to HYP CM + Man-GNPs (9.5 ± 2.4 × 10^4^, Fig. 4C). The GFAP stained volume did not change, but the ramifications emerging from astrocytic soma appeared significantly thicker in co-cultures exposed to HYP CM (average thickness 6.19 µm) compared to those exposed to NORM CM (4.33) or HYP CM + Man-GNPs (4.02, Fig. 4D). Ramification width was calculated with the full width at half maximum (FWHM) method. Microglia in the co-cultures were analyzed for their morphology. Iba1-stained microglia showed the sprouting of new ramifications after HYP CM exposure compared to NORM CM (Fig. 4E), indicative of their reactivity. This latter was counteracted in co-cultures exposed to HYP CM + Man-GNPs. Microglia signal was skeletonized for branch analysis, showing increased number of branches (11.5 ± 8.1, mean ± SD) and junctions (5.3 ± 4.1) when exposed to HYP CM compared to NORM CM (respectively, 3.6 ± 2.9 and 1.3 ± 1.5) and HYP CM + Man GNPs (4.8 ± 4.4 and 1.9 ± 2.4, Fig. 4F). Frequency distribution of each parameter is depicted in Fig. 4G.

**Figure 4.**
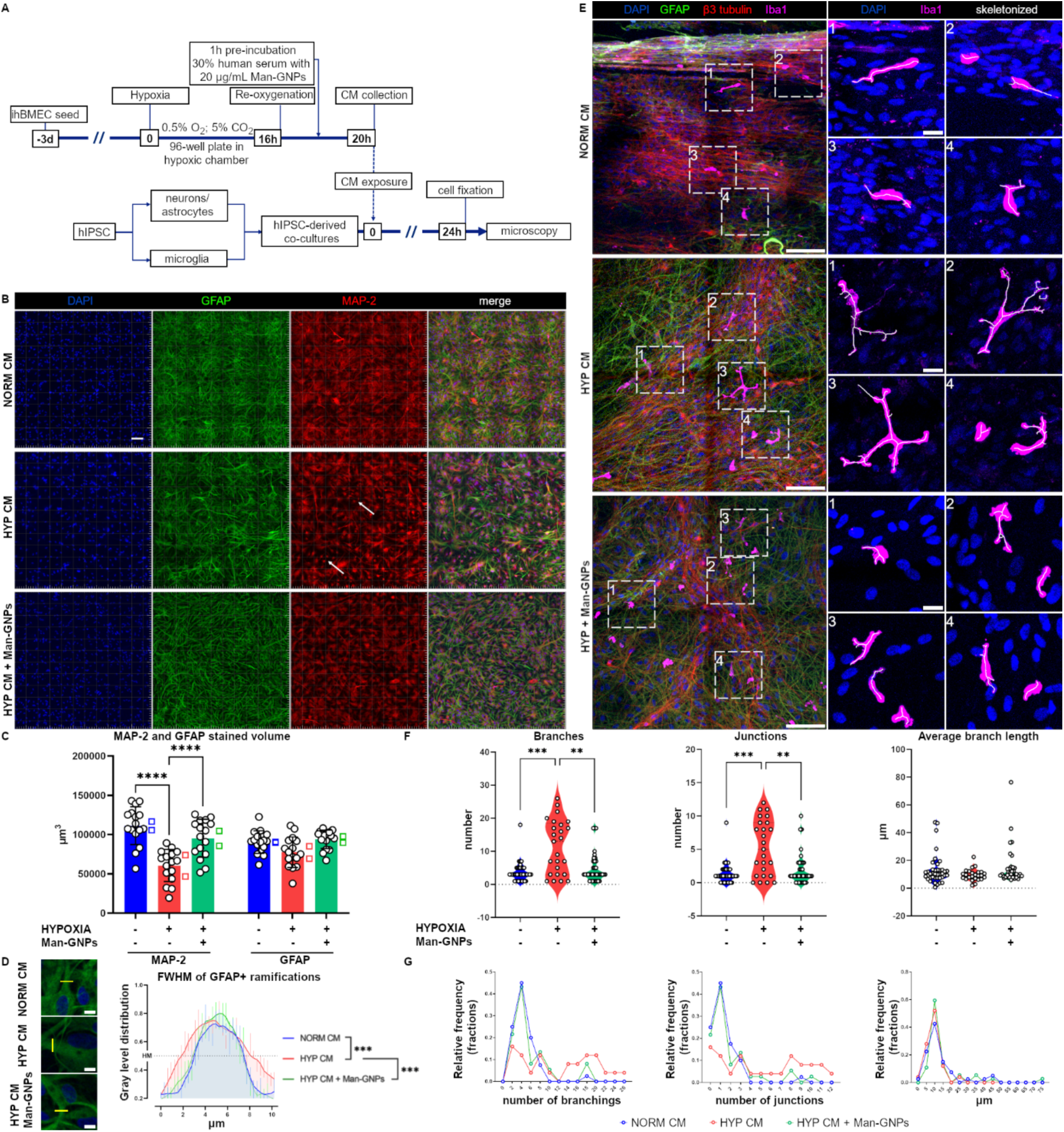
In vitro test of MBL inhibition by Man-GNPs, effects of ihBMECs’ conditioned medium on neurons, astrocytes and microglia. **A)** The experimental plan to generate ihBMECs’ normoxic or hypoxic conditioned medium (NORM CM, HYP CM, respectively) and co-cultures of hIPSC-derived neurons, astrocytes and microglia. **B**) Microphotographs of GFAP (astrocytes, green), MAP-2 (neurons, red) exposed for 24h to NORM CM or HYP CM +/− Man-GNPs. White arrows point to damaged neurons, i.e. circular cells without dendrites. Nuclei in blue (DAPI), scale bar 10 µm. **C**) The quantification of stained volumes (in µm^3^) showed a decrease of MAP-2 volumes in co-cultures exposed to HYP CM, which was counteracted by Man-GNPs. Data as mean ± SD. Each value is a random field of view (FOV) selected automatically from the overview image. Two-way ANOVA for repeated measures followed by Sidak’s multiple comparisons, ****p<0.0001 (n= 16 FOVs from two experimental replicates, empty rectangles indicate the mean of each replicate). **D**) Microphotographs of GFAP (green) and nuclei (DAPI, blue) with a yellow line along which we calculated the FWHM reported in the graph. Width of the first ramification emerging from astrocytic soma was calculated at gray level’s half maximum (HM) and was larger in HYP CM compared to NORM CM or HYP CM + Man-GNPs. Data as mean gray levels of 8 cells per group ± SEM. Two-way ANOVA followed by Tukey’s multiple comparisons, ***p<0.001. Scale bars 10 µm. **E**) Microphotographs of GFAP (green), β3-tubulin (neurons, red) and Iba1 (microglia, purple) exposed for 24h to NORM CM or HYP CM +/− Man-GNPs. Dashed squares indicate the magnified views of microglia on the right panels. Nuclei in blue (DAPI), scale bar 100 µm in full images, 20 µm in magnifications. White traces in the magnifications correspond to the Iba1 skeletonized signal. **F**) The quantification of microglia morphological parameters showed increased number of branches and junctions after HYP CM exposure, which was counteracted by Man-GNPs. Data as violin plot. Each dot is individual microglia. Kruskal-Wallis test, **p<0.01, ***p<0.001 (n= 25-40 cells from 3 FOVs placed in one well). **G**) Histograms of frequency distributions of the morphological parameters in E, shown with automatically chosen bin size.

### Characterization of the humanized mouse subjected to a model of ischemic stroke

Since no data were available as regards the characterization of hMBL KI mice in the context of ischemic stroke, we compared at first these mice with WT mice after transient occlusion of the middle cerebral artery (tMCAo) according to the experimental plan in Fig. 5A. At the experimental endpoint (2d after tMCAo), both strains had less viable neurons in the ipsi-lateral striatum (WT: 1273 ± 659.7 neurons/mm^2^ ± SD; hMBL KI: 571 ± 235.7) and cortex (WT: 1667 ± 538.5, hMBL KI: 1262 ± 527.5) compared to the contra-lateral side striatum (WT: 3023 ± 576.2, hMBL KI: 2760 ± 484.5) or cortex (WT: 3185 ± 418.8, hMBL KI: 3135 ± 559.9, Fig. 5B). The ischemic volume was similar between the strains, namely 49.5 ± 14.4 mm^3^ ± SD in WT and 41.9 ± 13.4 in hMBL KI (Fig. 5C and Supplementary Fig. S3). *hMbl2* was overexpressed in the liver of hMBL KI ischemic mice at 2d after tMCAo compared to sham (3.2 ± 0.46 fold-change ± SD), as seen previously in WT ischemic mice^43^, with no or negligible expression of the murine *Mbl1* or *Mbl2* (Fig. 5D). Western blot in non-reducing condition run over plasma samples at -2d, 90 minutes and 2d showed different MBL-A and MBL-C oligomerizations in WT mice (Fig. 5E, F, with raw blots presented in Supplementary Fig. S4). The total MBL-A and -C decreased at 2d compared to baseline -2d, as an index of protein consumption, i.e. binding to target. The relative abundance of specific oligomers in plasma changed over time, with 3×3 helix (hel), thus trimers of trimers of MBL-A and 2×3 hel (dimers of trimers) and monomers of MBL-C showing the most of changes on a time depended manner. In hMBL KI mice, plasma hMBL decreased at 2d indicating its consumption similarly to WT ischemic mice, but no clear-cut changes in the relative abundance of its oligomeric forms was observed over time (Fig. 5G). Fluorescence signals from hMBL merged with mouse-MASP-2 between 117-171 kDa (mMASP-2 in Fig. 5H). The plasma levels of mMASP-2 decreased over time similarly to those of hMBL (Fig. 5H, with raw blots presented in Supplementary Fig. S5), overall suggesting that hMBL was complexed with mMASP-2, and that their consumption could indicate LP activation.

**Figure 5.**
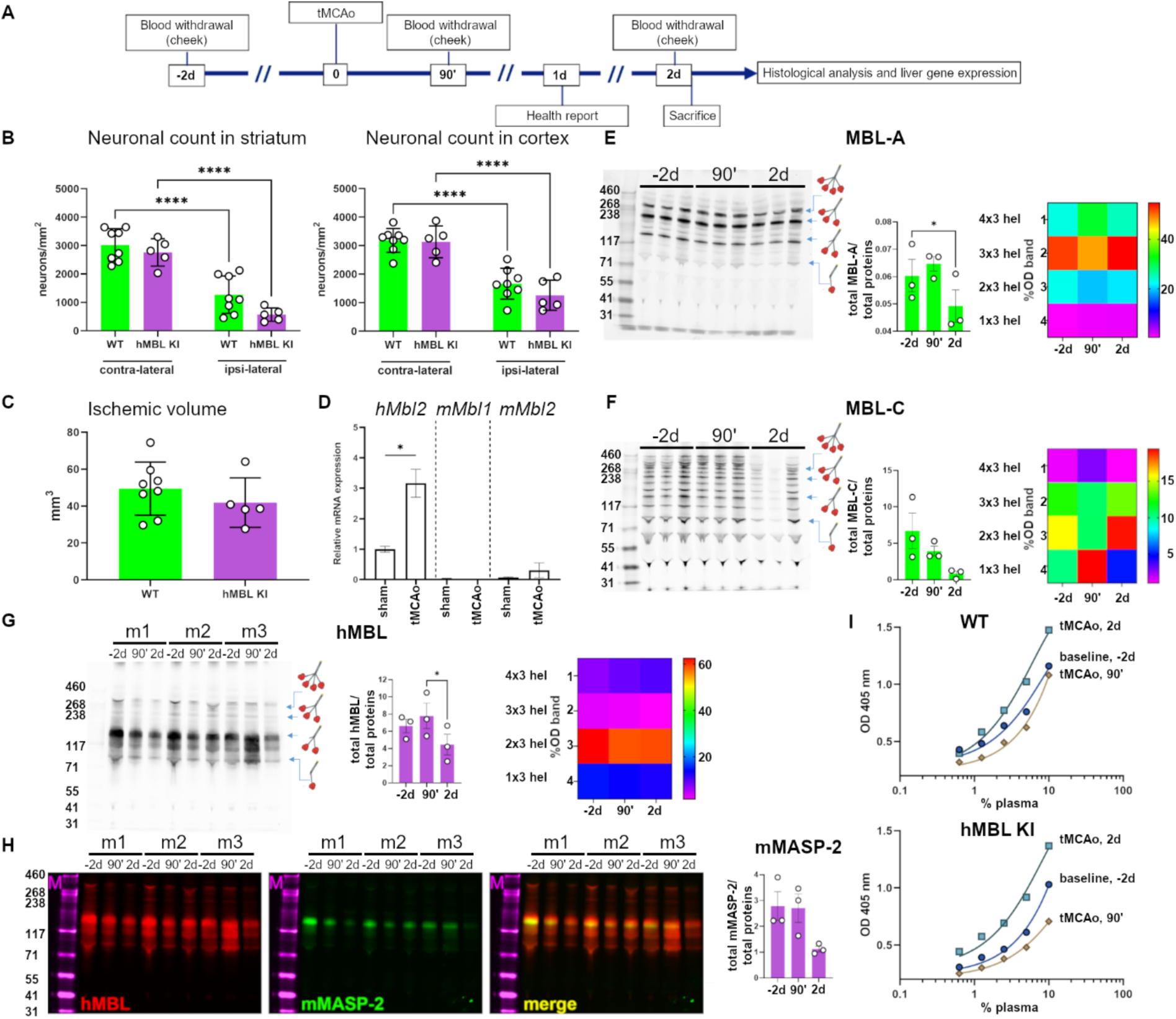
Characterization of the lectin pathway activation in a humanized mouse model of ischemic stroke. **A**) The experimental plan for the histological and biochemical analysis of hMBL knock in (hBML KI) mice subjected to the transient occlusion of the middle cerebral artery (tMCAo). **B)** At 2d after tMCAo, the number of neurons in striatum and cortex ipsi-lateral to the occlusion was reduced compared to the contra-lateral side in both WT and hMBL KI. Data as neurons/mm^3^ ± SD. Two-way RM ANOVA followed by Sidak’s multiple comparisons, ****p<0.0001 (n= 5-8). **C**) At 2d, the ischemic volume was similar in the two genotypes. Data as mm^3^ ± SD. Unpaired t-test (n= 5-8). **D**) At 2d, in the hMBL KI mice, *hMbl2* coding for hMBL was overexpressed in tMCAo compared to sham mice, with no expression of murine *mMbl1* and *mMbl2* isoforms. Data as bar ± SD, Unpaired t-test, *p<0.05. **E, F**) Western blot analysis for murine MBL-A (E) and MBL-C (F) in non-reducing condition over plasma samples of WT mice collected 2 days before (-2d) and 90 min (90’) and 48h after tMCAo. Total proteins decreased over time as a consequence of their consumption (i.e. target binding). Data as mean ± SD, One-way RM ANOVA followed by Dunnett’s multiple comparisons, *p<0.05 (n= 3). The color-coded heat map shows the relative abundance of the different oligomerized forms of MBL. **G**) Western blot analysis for hMBL in non-reducing condition over plasma samples of hMBL KI mice collected 2 days before (-2d) and 90 min (90’) and 48h after tMCAo. Total proteins decreased over time as a consequence of their consumption (i.e. target binding). Data as mean ± SD, One-way RM ANOVA followed by Dunnett’s multiple comparisons, *p<0.05 (n= 3). The color-coded heat map shows the relative abundance of the different oligomerized forms of MBL. **H**) The Western blot analysis in G with hMBL (red) and mMASP-2 (green) showing merged signal (yellow) between 117and 171 kDa. **I**) LP activity assay measuring C3b deposition on mannan-coated plates, showing similar activation in plasma from WT and hMBL KI mice. The data refer to pools of plasma from 5 mice per group, and plasma concentrations are reported on a logarithmic scale.

We then did an in vitro functional assay incubating mouse plasma (pools from 5 mice belonging to the same experimental group) on mannan-coated plates^43,54^, and measured the presence of C3b complement fragments. Both wild type and hMBL KI mice had a similar pattern of LP activation i.e., slight reduction at 90 minutes and increase at 2d after tMCAo compared to baseline plasma samples at -2d (Fig. 5I).

### In vivo glycoprofiling of brain ischemic vessels

Cryostate-cut brain sections of WT and hMBL KI ischemic mice were stained with fluolabelled WGA, ConA and IB4. The fluorescent signal from the lectins appeared to be increased in the ipsi-lateral cortex of WT and hMBL KI as compared to the contra-lateral cortex (Fig. 6A, A’). The quantification of the stained signal could show an increase for ConA – whose main target is α-D-mannosyl, in the ipsi-lateral cortex of WT (7.96 ± 2.60 × 10^8^ integrated density ± SD) and hMBL KI (8.22 ± 4.13 × 10^8^) compared to the contra-lateral side (WT: 3.98 ± 2.01 × 10^8^, hMBL KI: 3.76 ± 1.74 × 10^8^, Fig. 6B). WGA and IB4 signals increased similarly in the ischemic side, although, for hMBL KI, this was not statistically significant. The presence of hMBL was then assessed in ischemic hMBL KI mice, observing it on the vessels pertinent to the ischemic cortex, but not in those of the contra-lateral side (Fig. 6C, C’). The signal of hMBL was overlaid with that of fibrin in structures resembling clots before a vascular thinning, as indicated in Fig. 6D.

**Figure 6.**
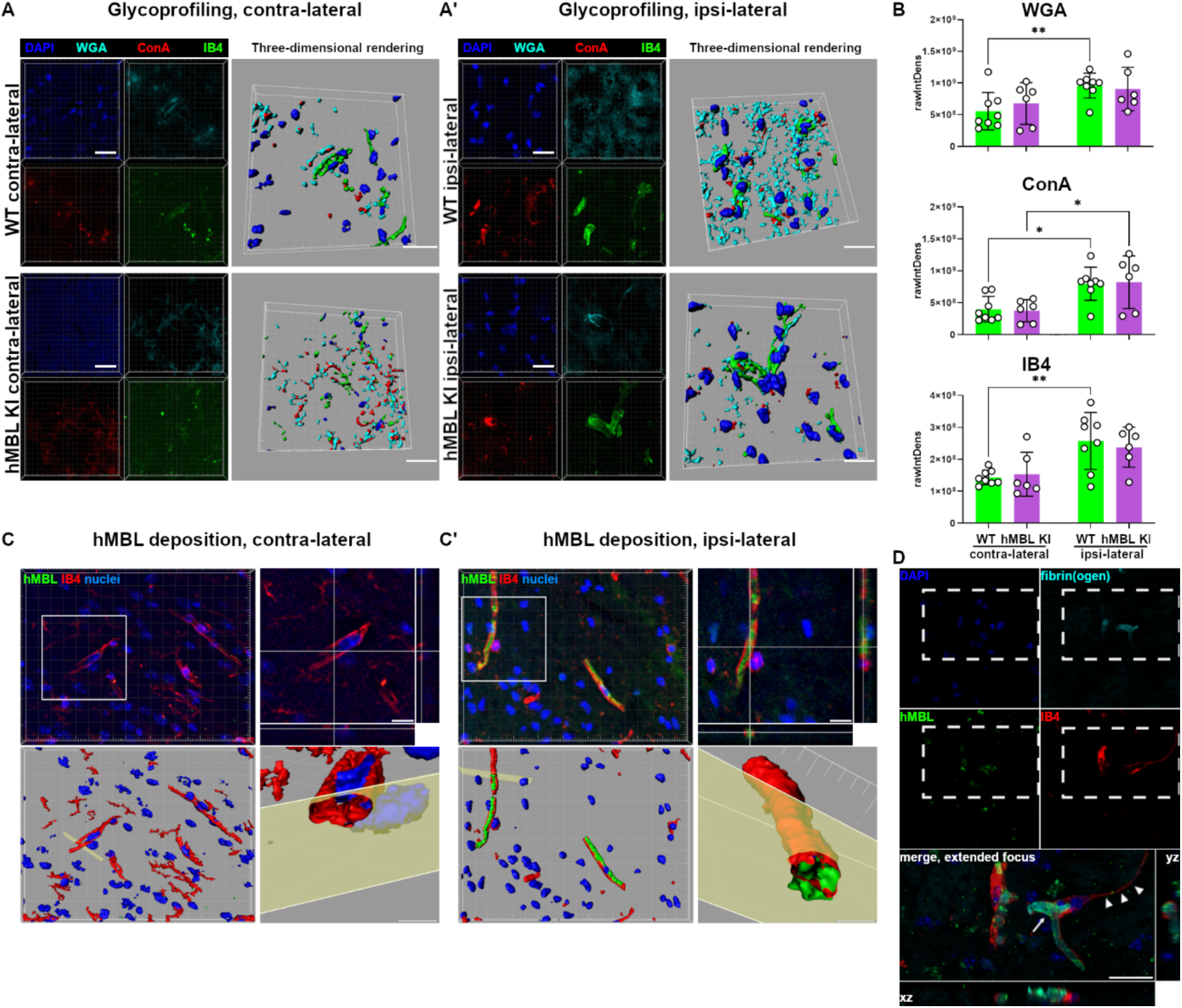
In vivo characterization of the brain vascular glycocalyx in a mouse model of ischemic stroke. **A, A’**) Microphotographs with 3D rendering of WGA (light blue), ConA (red) and IB4 (green) in contra-(A) and ipsi-lateral (A’) cortices of WT or hMBL KI mice at 2d after tMCAo. Nuclei in blue (DAPI), scale bars 20 µm. **B**) Quantification of the fluorescent signal intensity linked to the three plant lectins, showing increased WGA, ConA and IB4 in WT ipsi-lateral side compared to the contra-lateral and a similar trend for hMBL KI mice, which was statistically significant for ConA (labelling of α-D-mannosyl). Data as mean ± SD, Two-way ANOVA for repeated measures followed by Sidak’s multiple comparisons, **p<0.05, **p<0.001 (n= 6-8). **C, C’**) Microphotographs of hMBL (green) and vessels (IB4, red) in hMBL KI mouse at 2d after tMCAo. Single xy plane views with z projections of the white boxes in C and C’ confirm that hMBL was located in the luminal space of vessels. 3D renderings and clipped volumes (clipping in correspondence of the yellow planes) further demonstrate the intraluminal deposition of hMBL. No hMBL was detected on the vessels of healthy contra-lateral side. Nuclei in blue (DAPI), scale bars 10 μm. **D**) Microphotographs of co-localized fibrin (light blue) and hMBL (green) within an ischemic vessel (IB4, red). The co-localized signal could indicate a thrombus (arrow in the magnified area of the dashed rectangle) with downstream formation of string vessel (arrowheads). Nuclei in blue (DAPI), scale bar 20 µm.

### In vivo testing of Man-GNPs efficacy in ameliorating the outcome of ischemic hMBL KI mice

The last part of the study was aimed at evaluating if the promising properties of Man-GNPs, highlighted by in vitro binding studies and by functional cell assays, were maintained in vivo. For the sake of study’s translational value, we used a humanized mouse model (hMBL KI, both sexes), carrying the human MBL (encoded by *hMbl2*) and deleted of the two murine MBL isoforms^55^.

Male and female hMBL KI mice undergone tMCAo were treated with Man-GNPs, Glc GNPs or vehicle, administered intravenously 3h after the ischemic onset. Glyco-GNPs were used at a dose allowing to reach a concentration of 20 µg/mL in blood. Mice were followed according the IMPROVE guidelines over 8d of experimental endpoint (experimental plan in Fig. 7A). We observed a 25% mortality at the experimental endpoint for all experimental groups (Supplementary Fig. S6). GNPs-treated mice showed a slight and not significant decrease of sensorimotor deficits calculated as neuroscore^56^ at 2d after tMCAo (Fig. 7B). Man-GNPs-treated mice had a significantly reduced disinhibition-like behavior at the elevated plus maze (EPM, performed at 8d), as shown by increased difference of time spent in the closed-open arms (125.3 ± 94.9 seconds ± SD) compared to vehicle-treated (10.2 ± 138.1, Fig. 7C). The histological quantification of viable neurons at 8d after tMCAo showed improved neuronal survival in the ischemic cortex of Man-GNPs-treated mice (0.97 ± 0.18 ipsi-to-contra ratio ± SD) compared to vehicle treated (0.77 ± 0.15, Fig. 7D, D’ and E). We defined a combined outcome based on the quartile distribution of each assessed outcome - namely the neuroscore, the EPM and the neuronal quantification – finding that Man-GNPs treated mice could develop a good outcome more often than vehicle treated mice (Fig. 7F). The mice treated with Glc-GNPs could also have deficit attenuation, but to a lesser extent than Man-GNPs-treated. This latter observation is supported by the calculation of the effect of the treatment per their Odds ratio (OR) of association with a good outcome, showing, for Man-GNPs an OR 12.25 [1.52-81.92 CI 95%], for Glc-GNPs an OR 5.83 [0.76-38.05], Fig. 7G.

**Figure 7.**
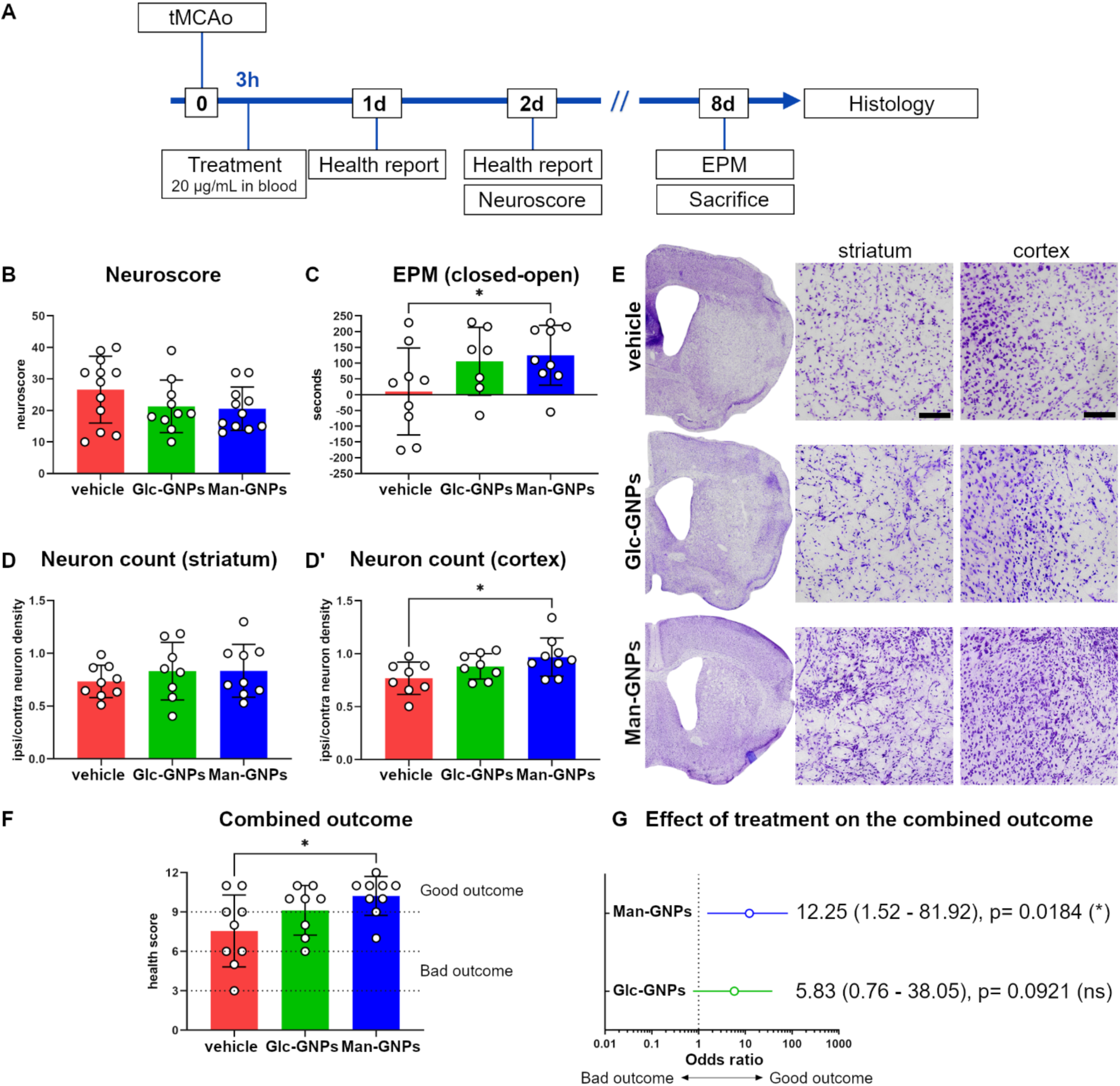
In vivo test of hMBL inhibition by Man-GNPs, effects on the outcome of hMBL KI tMCAo mice. **A**) The experimental plan to assess the ischemic outcome after treatment with Glc-GNPs or Man-GNPs 3h after the ischemic onset, using male and female hMBL KI tMCAo mice. **B**) Sensorimotor deficits by the neuroscore. **C**) Disinhibition-like behavior of mice assessed by the EPM at 8d after tMCAo. **D, D’**) Neuronal count in striatum (D) and cortex (D’) expressed as ratio of density in ipsi-. vs. contra-lateral side. Data in B-D panels as mean ± SD, One-way ANOVA followed by Fisher’s multiple comparisons, *p<0.05 (n=8-12, 1 mouse removed from Glc-GNPs in C as it did not performed the test). **E**) Microphotographs of Cresyl violet-stained brain sections as used for the neuronal count, scale bar 100 µm. **F**) A combined outcome was calculated using the behavioral and histological parameters. Mice receiving Man-GNPs had a score > 9 indicating a good outcome more frequently than the other groups. Data as mean ± SD, One-way ANOVA followed by Dunnett’s multiple comparisons, *p<0.05 (n=9). **G**) The odds ratio was calculated by a Chi square test using the Baptista-Pike method for calculating the 95% confidence interval (CI 95%), stratifying mice in terms of good outcome (defined as a score > 9) versus bad outcome (score ≤ 6). Forest plot showing that the Man-GNP treatment was significantly associated with a good outcome.

## Discussion

The present data show that Man-GNPs prevent MBL’s deposit on its glycoproteic targets on ischemic brain microvascular endothelial cells, and the consequent activation of inflammatory cascades. Man-GNPs also demonstrated a relatively small, but significant, protection in vivo, in a hMBL KI mouse model of ischemic stroke. That MBL is pathogenic in acute brain injury has been previously reported by us and others^57–60^. Here, we provide experimental support to our early working hypothesis that modified glycoproteins on the ischemic vessels offer binding targets to MBL. We found that the ischemic vessels show a higher binding of plant lectins specific for α-D-mannosyl (ConA) or N-acetylglucosaminyl (WGA), which are the main targets of MBL. Both glycans are components of the glycocalyx of endothelial cells, the inner (i.e. luminal) component of the tripartite BBB^5^, which undergoes critical modification after acute injury preceding the activation of thrombo-inflammatory pathways^39^. Based on literature data, an initial release of glycans from the damaged endothelium is regarded as a source of DAMPs, as well as of reduced BBB function^3^. Previous studies of ischemia/reperfusion injury in renal^61^, cardiac^62^, hepatic^63^, and cerebral tissues^64,65^ have assessed glycocalyx damage indirectly by detecting proteoglycans or related enzymes released into the circulation. In contrast, we directly quantified specific sugar moieties on injured endothelial cells as source of brain local DAMPs for MBL. While the endothelial damage evolves, glycocalyx changes are likely to be specific for the vessel type and the diseased condition, but have not been fully explored yet. In our hypoxia/re-oxygenation model using brain microvascular cells, we observed a loss of glycan targeting by lectins soon after hypoxia, followed by increased lectin binding after re-oxygenation. It remains to be understood whether the target glycans of MBL become available through their new synthesis or favored accessibility. Regardless, MBL deposition was selectively observed in vessels pertinent to the ischemic territory after re-oxygenation. Being a liver-synthetized circulating protein, intercepting MBL in the blood before its vascular deposition should provide protection.

It is important to recall that MBL has an oligomeric structure with multiple carbohydrate recognition domains (CRD), allowing multiple interactions with arrays of polyglycosylated targets with high avidity^8,41^. Thus, to effectively interfere with these multivalent interactions, multivalent inhibitors such polymannosylated compounds^27,28,66^ are required to fully leverage this multivalency effect. In this regard, nanoparticles can provide a mean to increase the multivalent binding of MBL, as previously demonstrated with Man-GNPs, achieving MBL binding to at least 2–3 orders of magnitude greater than monomeric mannose^42^. Building on these findings, in this work, we further explored the use of these Man-GNPs to better understand their potential physiological effects. Nanoparticles represent an important application of nanobiotechnology to medicine and there are already many commercially available examples, including Copaxone for multiple sclerosis, Zevalin for lymphoma and Abraxane for cancer^67,68^. Their unique properties, and in particular the possibility to functionalize them with labels or ligands, allow their applications for molecular diagnostics and targeted drug delivery, with the aim of avoiding side effects and improving the therapeutic window ^69,70^. After demonstrating the lack of in vitro toxicity of glyco-GNPs, we showed that 20 µg/mL Man-GNPs significantly reduced MBL deposition on the hypoxic ihBMECs after re-oxygenation, an effect that was associated to a significant reduction of the hypoxia-induced over-expression of the inflammatory gene *ICAM-1* (*MMP-2* was also reduced to a similar extent, although the decrease was not statistically significant). *ICAM-1* encodes for a transmembrane protein expressed by activated endothelial cells and is involved in the recruitment of inflammatory cells to the injured brain, i.e. facilitates leukocyte endothelial transmigration^71^. Indeed the connection of ICAM-1 and MBL was shown before in vivo as ICAM-1 expression is downstream to the MBL deposition on ischemic endothelium^39^. The results obtained here with the in vitro cellular model of hypoxic ihBMEC are thus consistent with these previous in vivo results. No inhibitory effect of Man-GNPs was observed at the highest dose tested here (40 µg/mL). It may be speculated that this was due to aggregation of the nanoparticles at this concentration, thus lowering their available functionalized surface to MBL binding.

To clarify if Man-GNPs may target MBL in the cells, co-localization studies were carried out. The binding of Man-GNPs to MBL, and thus the above-mentioned effects due to MBL scavenging, are likely to occur in the extracellular space, since no co-localization of the two was observed at the intracellular level. This aligns with MBL binding to cells via its CRD, which also serves as the binding site for Man-GNPs. Consequently, after MBL is deposited on the cells, the CRD may no longer be accessible to Man-GNPs.

We next studied in vitro the consequences of MBL vascular deposition and of its inhibition on neuronal viability and glial cells activation. For this, we collected the conditioned media (CM) from the ihBMEC experiments and applied them to human co-cultures of neurons, astrocytes and microglia, all differentiated from hIPSC from a healthy donor. The hypoxic CM induced neuronal damage, measured as reduced MAP-2-stained volume, and neuroinflammation, this latter seen as thicker astrocytic branches and increased microglia ramifications compared to the normoxic CM. Notably, neuronal damage and neuroinflammation were reduced when the hypoxic CM was collected from Man-GNPs-treated ihBMEC. These data suggest that the vascular protective effect of our MBL-targeting strategy may mitigate the harmful effects of hypoxia on the neural environment.

The complement system is highly preserved evolutionally, but, likely due to its protein redundancy, it shows species-selective differences in initiators and effector molecules. In rodents two functional MBL isoforms (-A and -C) are present, at variance with the only human protein.

It has been proposed that MBL genes duplicated prior to human-rodent divergence, and that the human homolog to *Mbl1* (coding for MBL-A) was lost during evolution^72^. Murine MBL isoforms differ in monosaccharide specificity, in their polymerization state and hence they may have different accessibility to microorganisms and DAMPs^73^. In brain ischemia we previously reported an earlier MBL-A vascular deposition than MBL-C, which implied a larger contribution of the former isoform to the ischemic injury^43^. The human MBL shares carbohydrate specificity with MBL-C, in agreement with the high homogeneity between the murine and the human *Mbl2* gene^72,73^. Although it is MBL-A (less similar to human MBL) to be the key contributor to brain ischemic injury, clinical data show that stroke patients with MBL deficiency develop less severe injury^57^, thus indicating human MBL as a pharmacological target.

Since our glyco-GNPs were developed and tested in vitro against the human MBL, we explored their therapeutic potential in a mouse line carrying the human protein, but not the murine isoforms. The hMBL KI mice were subjected to ischemic stroke, induced by transient (30 min) occlusion of the middle cerebral artery (tMCAo) followed by reperfusion. Of note, the humanized mice were only used in kidney ischemia, thus requiring a preliminary characterization in the context of ischemic stroke. It was found that hMBL showed a similar oligomerization before and after tMCAo, which is important for more reliable translation of the results since in vitro affinity studies of Man-GNPs were carried on serum samples collected from healthy donors^42^. Indeed, hMBL oligomerization states may alter the availability of their carbohydrate binding sites to the Man-GNPs. Moreover, our data suggest that circulating hMBL is associated with MASP-2 (mannose-binding lectin-associated serine protease-2) - the first enzymatic component for the lectin pathway of complement activation^74,75^. As observed in WT mice, analysis of the ischemic vessels in the humanized strain revealed that ischemia induced the exposure of mannose-containing DAMPs (detected using fluorescent ConA), as well as MBL deposition. This was accompanied by activation of the LP, as indicated by in vitro measurements of C3b formation in plasma before and after ischemia, and co-localization with fibrin.

After the characterization of the hMBL KI mouse model of ischemia, reporting the relevance of MBL and the lectin pathway in the ischemic stroke model, we evaluated if the consequences of ischemia could be ameliorated by Man-GNPs, administered systemically (intravenously) 3h after reperfusion, at dosing yielding a plasma concentration of 20 µg/mL. Man-GNPs induced a small, non-significant improvement of sensorimotor deficits at 48h after tMCAo, while a larger protection was seen at 8d. At this latter time point, Man-GNPs-treated mice performed better at the elevated plus maze, and displayed reduced neuronal loss compared to vehicle-treated. We also analysed a group of mice administered with Glc-GNPs, which bind MBL with >25-fold lower affinity than Man-GNPs as per previous SPR characterization^42^. Glc-GNPs-treated mice showed a tendency in ischemic deficit amelioration, which may suggest MBL (or other lectins) targeting in vivo. This is in line with the observation that MBL was present on the soft protein corona of Glc-GNPs, which may provide a sufficient inhibitory effect in vivo. However, overall, our glyco-GNPs did not provide a wide effect size on ischemic protection, as observed with other inhibitory compounds^27,76^. We may hypothesize a rapid removal of glyco-GNPs from the circulation for a very fast elimination and/or accumulation in organs^77–79^ (i.e. liver, kidney, spleen), thus reducing their bioavailability in blood. Further pharmacokinetic studies are needed to clarify this point as well as to define a more effective treatment schedule, including multiple dosing. Another relevant point is the use of male and female mice in our in vivo experiment. This is a highly recommended improvement in the use of the ischemic model, but may introduce some outcome variability. Namely, we observed reduced deficits in vehicle-treated female compared to male mice, with these latter having the best protective effect when administered with Man-GNPs. However, the use of both sexes of a humanized mouse model favors the solidity and generalizability of our data, supporting the pharmacological value of multivalent glyco-GNPs to target inflammatory lectins after acute brain injury.

## Acknowledgements

Pancaro Alessia and Nelissen Inge from the Flemish Institute for Technological Research (VITO nv), Health Unit, Mol, Belgium.

Africa Garcia Barrientos and John Porter from Midatech Pharma PLC, Cardiff, UK.

## Author contributions

GM, MG, SF planned and executed the experiments, analyzed the data and drafted the ms DM, M-GDS designed the work, analyzed the data, drafted the ms

SS, AB, AP, CD, PPS, LP, CD, FO, LF, DC, GF, MDP executed the experiments and analyzed the data

