## Supplementary Figures for "Glycan-coated nanoparticles mimicking the ischemic glycocalyx scavenge the complement system conferring protection after experimental ischemic stroke"

**Supplementary Material**

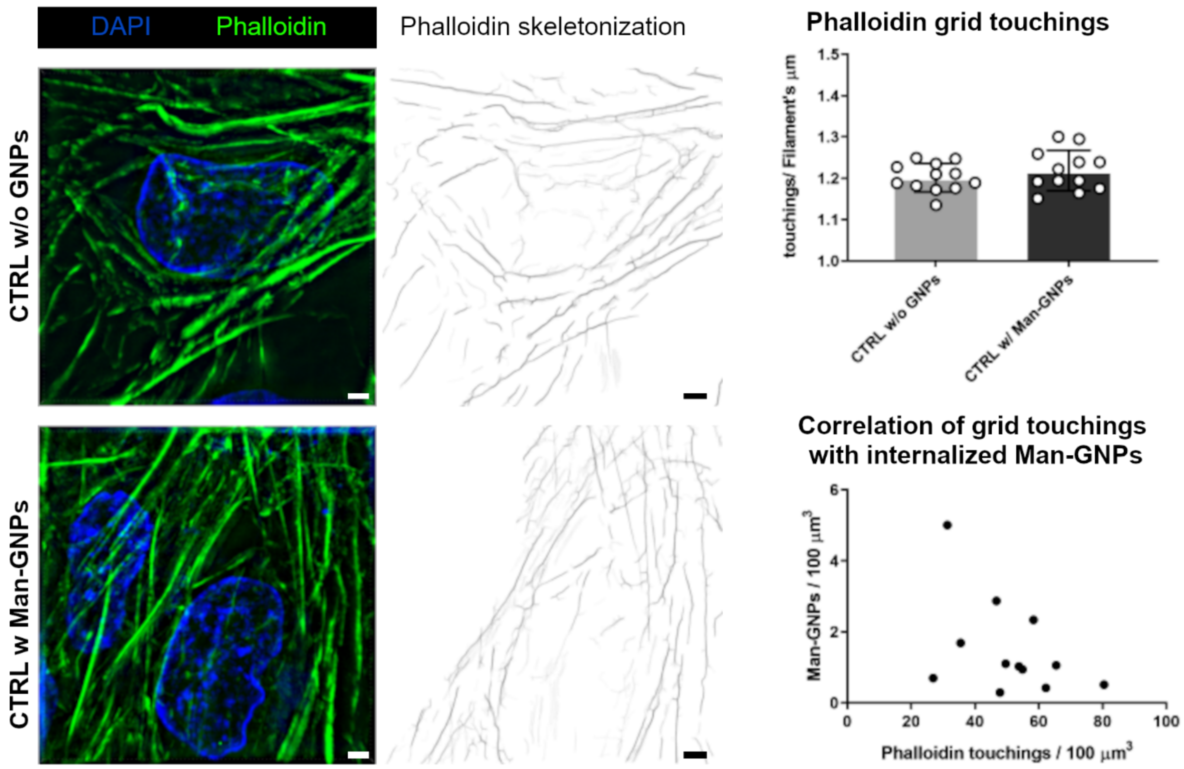


**Supplementary Figure S1.** Test of Man-GNPs in vitro toxicity.

To investigate possible toxicity of Man-GNPs, normoxic ihBMECs (CTRL) exposed to them were randomly selected and analysed for structural changes of their cytoskeleton compared to unexposed ihBMECs. F-actin filaments labelled by phalloidin were visualized by superresolved structured illumination microscopy (SIM) and quantified by recording their touching with a superimposed grid. The panels show SIM images of F-actin (phalloidin, green) were skeletonized and a 1 mm-spaced grid was superimposed. Nuclei in blue (DAPI), scale bars 2 µm. Touching per filament length unit (1 µm) in Man-GNPs-exposed ihBMEC did not differ compared to those unexposed, and no correlation was found between the touching and Man-GNPs number per 100 $\mu m^3$ of analyzed area. Data as mean with individual values ± SD (n=12 cells from four wells per condition). Unpaired t test, ns.

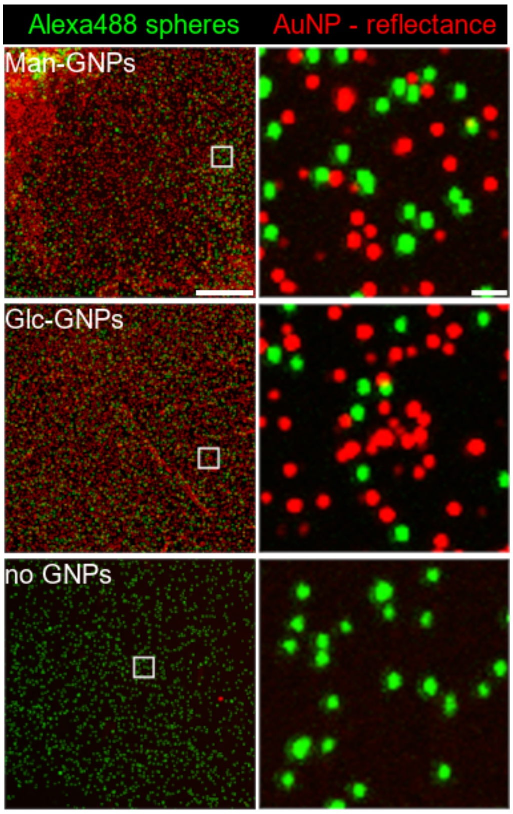


**Supplementary Figure S2.** Set up of GNP visualization using reflectance confocal microscopy.

The presence of gold quenches fluorescent dyes, hence the GNPs could not be visualized through fluorescent tags. Thus, we visualized the GNPs by reflectance confocal microscopy (RCM). First, RCM was set up by illuminating the metallic surface of glyco-GNPs spotted on a microscopy glass with a single-wavelength laser (561 nm) and then collecting the reflective light at the same wavelength. The focus was adjusted by Alexa488-conjugated fluorescence beads which were added to the same spot of the GNPs. The panels show microphotographs of sugar-GNPs (red) spotted on glasses and captured by reflectance confocal microscopy (the white squares indicate the magnified field of view). Focus adjustment was done by Alexa488-conjugated beads (100 nm, green). Control images only in presence of Alexa488-conjugated beads presented as no GNPs. Scale bars 20 µm in full image, 1 µm in magnification.

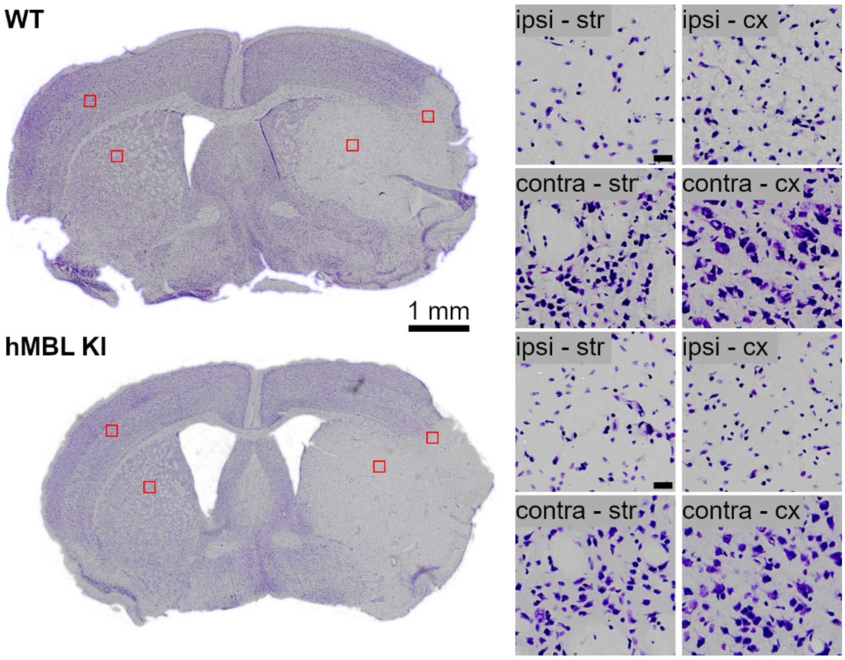


**Supplementary Figure S3.** Cresyl violet-stained sections.

Microphotographs of Cresyl violet-stained brain sections of WT (upper panels) or hMBL KI (lower) as used for the neuronal count. Magnified fields of ipsi- and contra-lateral striatum and cortex are positioned where indicated by the red boxes. Scale bar 1 mm in the whole sections, 20 µm in magnifications.

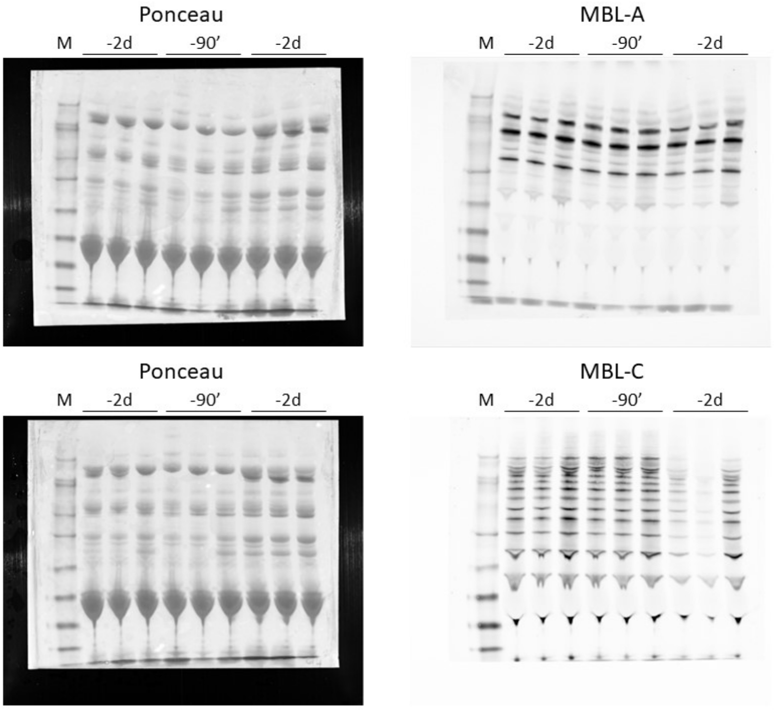


**Supplementary Figure S4.** Raw western blots images.

Western blot analysis for murine MBL-A (upper panels) and MBL-C (lower) in non-reducing condition over plasma samples of WT mice collected 2 days before (-2d), 90 min (90') and 48h after tMCAo (M stands for 'marker'). The Ponceau staining shows the total amount of proteins present in the analyzed plasmas and was used to normalize the densitometric quantification of each band of MBL-A or -C blots.

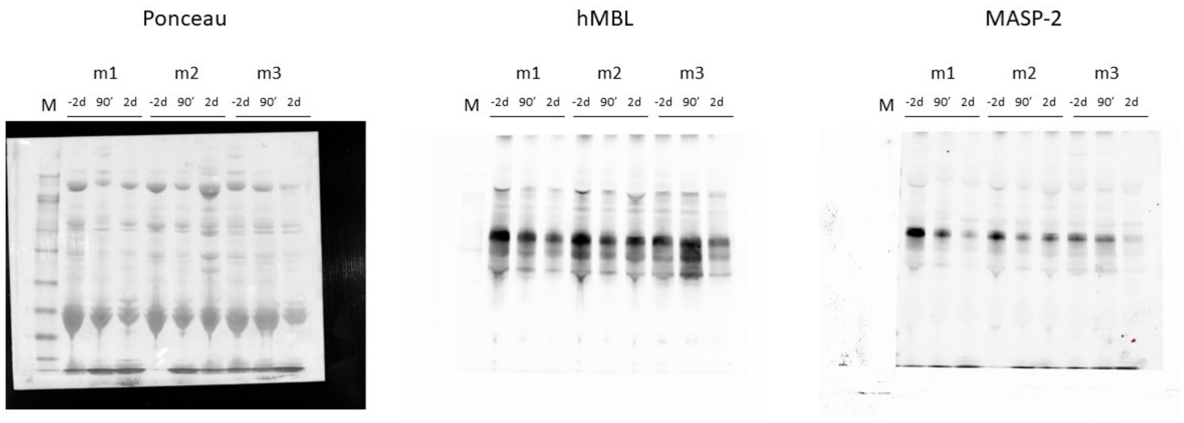


**Supplementary Figure S5.** Raw western blots images.

Western blot analysis for human MBL and murine MASP-2 in non-reducing condition over plasma samples of WT mice collected 2 days before (-2d), 90 min (90') and 48h after tMCAo (M stands for 'marker', m for each mouse analyzed longitudinally). The Ponceau staining shows the total amount of proteins present in the analyzed plasmas and was used to normalize the densitometric quantification of each band of hMBL and MASP-2 blots.

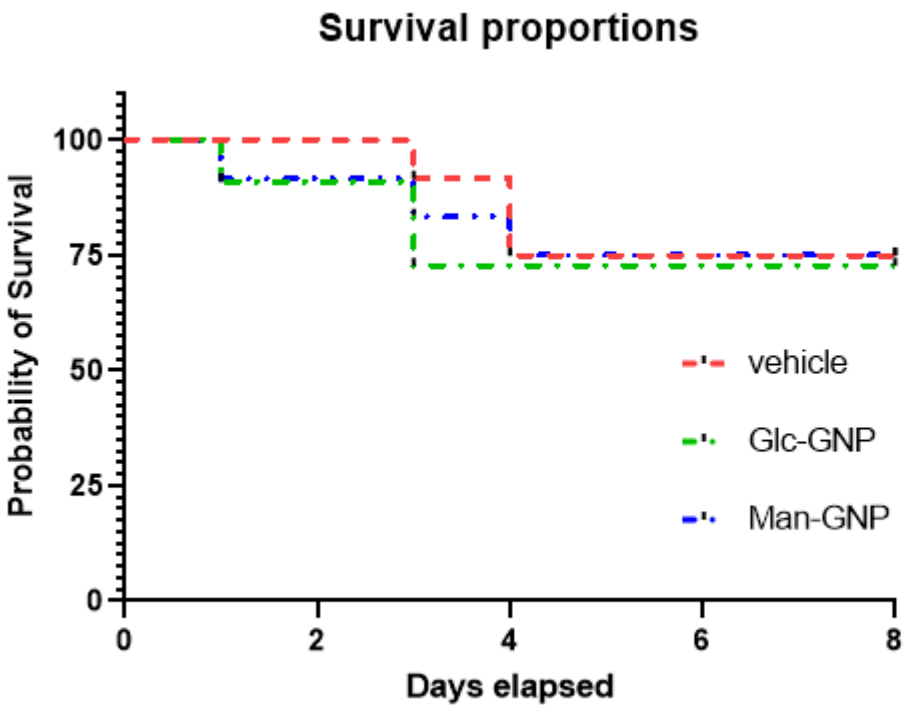


**Supplementary Figure S6.** Survival proportions of ischemic hMBL KI mice receiving vehicle or glyco-GNPs, within the experimental endpoint (8d after tMCAo).
